# Genome of the parasitoid wasp *Diachasma alloeum*, an emerging model for ecological speciation and transitions to asexual reproduction

**DOI:** 10.1101/545277

**Authors:** Eric S. Tvedte, Kimberly K.O. Walden, Kyle E. McElroy, John H. Werren, Andrew A. Forbes, Glen R. Hood, John M. Logsdon, Jeffrey L. Feder, Hugh M. Robertson

**Author notes:** Authors for Correspondence: Eric Tvedte, Institute for Genome Sciences, University of Maryland School of Medicine, Baltimore, MD, 21201. Hugh M. Robertson, Department of Entomology, University of Illinois at Urbana-Champaign, Urbana, IL, 61801. Data deposition: The Raw read data may be accessed on NCBI SRA database and can be accessed via BioProject portals for genome (PRJNA284396) and transcriptome (PRJNA283787) datasets.

## Abstract

Parasitoid wasps are among the most speciose animals, yet have relatively few available genomic resources. We report a draft genome assembly of the wasp *Diachasma alloeum* (Hymenoptera: Braconidae), a host-specific parasitoid of the apple maggot fly *Rhagoletis pomonella* (Diptera: Tephritidae) and a developing model for understanding how ecological speciation can “cascade” across trophic levels. Identification of gene content confirmed the overall quality of the draft genome, and we manually annotated ∼400 genes as part of this study, including those involved in oxidative phosphorylation, chemosensation, and reproduction. Through comparisons to model hymenopterans such as the European honeybee *Apis mellifera* and parasitoid wasp *Nasonia vitripennis*, as well as a more closely related braconid parasitoid *Microplitis demolitor*, we identified a proliferation of transposable elements in the genome, an expansion of chemosensory genes in *D. alloeum* and other parasitoid wasps, and the maintenance of several key genes with known roles in sexual reproduction and sex determination. The *D. alloeum* genome will provide a valuable resource for comparative genomics studies in Hymenoptera as well as specific investigations into the genomic changes associated with ecological speciation and transitions to asexuality.

## Introduction

The Hymenoptera may be the largest order of insects due to the immense diversity of parasitic wasps (*i.e.* “parasitoids”) that lay their eggs into or on other insect species (LaSalle & Gould, 1993; Austin & Dowton, 2000; Whitfield, 2003; Forbes *et al*., 2018). The great diversity of parasitoid wasps may be a consequence of their close relationship with their insect hosts. When a specialist parasitoid shifts to a new host, this change can propel the evolution of reproductive isolating barriers between wasp populations using the new and ancestral hosts (Feder & Forbes, 2010). The evolution of reproductive isolating barriers following a host shift is a well-documented phenomenon in host specialist insects (Forbes *et al*., 2017), but the study of genomic changes that accompany such phenomena is still in its early stages.

*Diachasma alloeum* (Hymenoptera: Braconidae) is a specialist parasitoid of the fruit fly *Rhagoletis pomonella* (Diptera: Tephritidae). After the introduction of domesticated apples to the United States from Europe, *R. pomonella* infesting native hawthorn fruits experienced a host shift and subsequently evolved reproductive isolating barriers in what has become a well-known example of incipient ecological speciation (Walsh, 1867; Bush, 1966; Bush, 1994; Nosil, 2012). This new “apple maggot fly” was sequentially colonized by *D. alloeum*, which appears to have shifted from its ancestral host, the blueberry maggot *Rhagoletis mendax* (Forbes *et al*., 2009). Two reproductive isolating barriers (*i.e.* diapause emergence and host fruit volatile discrimination) have evolved in parallel in *R. pomonella* and *D. alloeum*, and in both fly and wasp, these traits appear to have a genetic basis (Dambroski *et al*., 2005, Forbes & Feder, 2006, Forbes *et al*., 2009). This phenomenon of “sequential” or “cascading” speciation may be an important driver of new biodiversity (Stireman *et al*., 2006; Abrahamson & Blair, 2007; Hood *et al*., 2015).

Reproductive isolation in genus *Diachasma* has also arisen as a consequence of the loss of sexual reproduction, a general pattern observed in many hymenopteran insects (van der Kooi *et al*., 2017). Asexual *D. muliebre* appears to have split from its sexual sister *D. ferrugineum* between 0.5 and 1 mya (Wharton & Marsh, 1978; Forbes *et al*., 2013). Although the decay of genes involved in sexual traits has been observed in multiple asexual parasitoid wasps (*e.g.* Ma *et al*., 2014; Kraaijeveld *et al*., 2016), there is a dearth of comparative assessments of genomic molecular evolution between sexual and asexual Hymenoptera.

Here, we report the *de novo* genome assembly of the parasitoid wasp *D. alloeum*, adding to the genomic resources for parasitoid wasps, which are underrepresented among available hymenopteran genomes (Branstetter *et al*., 2017). We performed a series of descriptive analyses to assess the overall quality and content of the *D. alloeum* genome, and then focused on annotation and evolutionary analyses of gene families with potential relevance to speciation and sex determination in *Diachasma*.

## Results and Discussion

### Quality assessment of genome assembly

Libraries from a combination of single and pooled wasp samples contained 182.88 GB total sequence data. The *de novo* genome assembly Dall1.0 (GenBank accession: GCA_001412515.1) had 3,968 scaffolds with a total scaffold length of 388.8 Mb and a scaffold N50 of 645,583 bp (Table 1). The presence of prokaryotic-like sequences in eukaryotic genome projects may reflect contamination in sequencing libraries or an actual association between microorganisms and hosts. Of the *D. alloeum* scaffolds, we annotated 656 as likely bacterial contaminants and an additional scaffold (RefSeq accession: NW_015145431.1) as an apparent lateral gene transfer event from a *Rickettsia* species (Supplementary Material online). The likely bacterial contaminating scaffolds were removed from the *D. alloeum* assembly, and the remaining 3,313 scaffolds will be made available as genome assembly Dall2.0.

**Table 1.** Summary statistics and feature counts of *D. alloeum* genome assembly.

| Assembly | **Dall1.0** | **Dall2.0** |
| --- | --- | --- |
| Total Sequence Length | 388,752,668 | 384,371,871 |
| Scaffold Count | 3,968 | 3,313 |
| Scaffold N50 | 645,483 | 657,001 |
| Contig Count | 25,534 | 24,824 |
| Contig N50 | 44,932 | 45,492 |
| GC% | 38.45 | 38.30 |

A common metric used to assess the relative completeness of a genome assembly is the identification of conserved single-copy genes, performed here using BUSCO v3 (Simão *et al*., 2015). We found 1060/1066 (99%) arthropod-specific BUSCO genes in the *D. alloeum* genome, including 1053/1066 complete genes. These values are similar to BUSCO gene content in other published hymenopteran genomes (Table 2, Supplementary Material online). Our *de novo* assembly of the *D. alloeum* mitochondrial sequence using NOVOplasty (Dierckxsens *et al*., 2017) produced a 15,936 bp sequence with a complete set of thirteen protein coding genes, two rRNA sequences, and 20 tRNA sequences. In addition, our annotation of 65/68 (96%) of the canonical suite of nuclear-encoded mitochondrial genes provided additional evidence for a highquality genome assembly (Supplementary Material online).

**Table 2.** Summary statistics and BUSCO gene content for genome assemblies in four hymenopteran insects.

| Organism | Assembly Accession | Assembly Size | Scaffold Count | Scaffold N50 | Complete arthropod BUSCOs |
| --- | --- | --- | --- | --- | --- |
| *Diachasma alloeum* | GCA_001412515.1<br>(this study) | 388,752,668 | 3,698 | 645,483 | 1053 (99%) |
| *Apis mellifera* | GCA_000002195.1<br>(Elsik *et al.*, 2014) | 250,287,000 | 5,645 | 997,192 | 1044 (98%) |
| *Nasonia vitripennis* | GCA_000002325.2<br>(Werren *et al.*, 2010) | 295,780,872 | 6,169 | 708,988 | 1038 (97%) |
| *Microplitis demolitor* | GCA_000572035.2<br>(Burke *et al.*, 2018) | 241,190,213 | 1,794 | 1,139,389 | 1057 (99%) |

We used RepeatModeler (Smit *et al*., 2015), PASTEClassifier (Hoede *et al*., 2014, version 1.0) and RepeatMasker (Smit *et al*., 2010) for *de novo* repeat identification, repeat reclassification, and repeat quantification, respectively (Supplementary Material online). Remarkably, nearly half (49%) of the *D. alloeum* genome consisted of repetitive sequences, although a substantial contributor (30%) was from unclassified repetitive sequences.

### Expansion of species-specific chemosensory genes in *D. alloeum*

Chemoreception in arthropods is mediated by three major families of receptors: odorant receptors (**ORs**), gustatory receptors (**GRs**), and ionotropic receptors (**IRs**) (Clyne *et al*., 1999; Clyne *et al*. 2000; Benton *et al*., 2009). In addition, two major families of water-soluble proteins are responsible for transport and/or quenching of ligands to chemosensory receptors: odorant binding proteins (**OBPs**) and chemosensory proteins (**CSPs**) (Vieira & Rozas, 2011; Pelosi *et al*., 2014; Larter *et al*., 2016). Chemosensory discrimination of fruit volatiles is an important axis of divergence among host fly-associated populations of *D. alloeum*, initiating reproductive isolating barriers between these wasps (Forbes *et al*., 2009).

Previous characterizations of chemosensory genes in hymenopteran insects, in particular the gene-rich receptor families, demonstrate that automated gene prediction pipelines are generally poor at accurately predicting these gene models (Robertson & Wanner, 2006; Robertson *et al*., 2010; Croset *et al*., 2010; Zhou *et al*., 2015; Robertson *et al*., 2018). We therefore manually annotated a total of 321 gene models that represents the full inventory of five chemosensory gene families in *D. alloeum* (Table 3, Supplementary Material online). Consistent with GO-enrichment analyses produced by OrthoVenn, we found lineages of OR expansions in *D. alloeum*, and clusters of GR homologs present in the braconid wasps *D. alloeum* and *M. demolitor* but absent in the well-studied hymenopterans *Nasonia vitripennis* or *Apis mellifera*. We also observed a notable expansion of IRs in *D. alloeum* relative to another *Microplitis* species, *M. mediator* (Figure 2). As odor discrimination has likely contributed to reproductive isolation following a host shift in *D. alloeum*, this dataset will be important for the study of chemosensory gene composition and evolutionary rate differences within and between *Diachasma* species.

**Figure 1.**
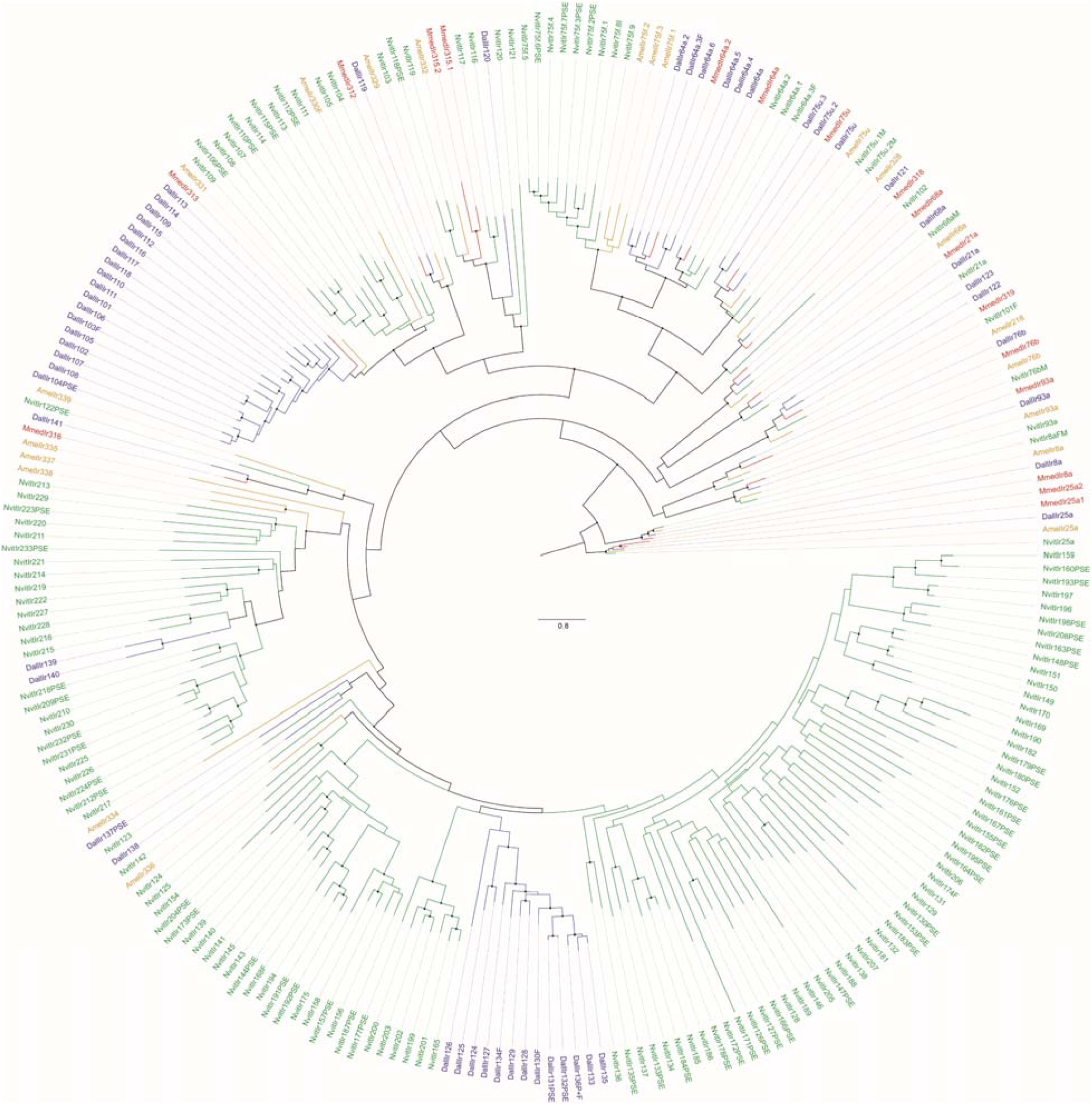
Phylogenetic tree of ionotropic receptors (IRs) from sampled hymenopteran insects. Dall = *Diachasma alloeum*, Mmed = *Microplitis mediator*, Nvit = *Nasonia vitripennis*, Amel = *Apis mellifera*. Maximum likelihood tree generated using 656 alignment columns. Dots on nodes indicate > 90% bootstrap support. The scale bar indicates the number of amino acid substitutions per site.

**Table 3.** Chemosensory gene content of selected hymenopteran insects.

| Organism | ORs | GRs | IRs | OBPs | CSPs | Citations |
| --- | --- | --- | --- | --- | --- | --- |
| *D. alloeum* | 187(14) | 39(1) | 51(5) | 15(0) | 9(0) | This study |
| *A. mellifera* | 163 (11) | 10(0) | 10(0<sup>a</sup>) | 21(0<sup>a</sup>) | 6(0<sup>a</sup>) | Robertson & Wanner, 2006<br>Foret & Maleszka, 2006<br>Foret *et al.*, 2007<br>Croset *et al.*, 2010 |
| *N. vitripennis* | 225 (76) | 47(11) | 99(54) | 90(8) | 9(0) | Robertson *et al.*, 2010<br>Robertson *et al.*, 2018<br>Werren *et al.*, 2010<br>Vieira *et al.*, 2012 |
| *M. demolitor*<sup>b</sup> | 218 (4) | 85(1) |  |  |  | Zhou *et al.*, 2015 |
| *M. mediator* |  |  | 17(0<sup>a</sup>) | 20(0<sup>a</sup>) | 3(0<sup>a</sup>) | Zhang *et al.*, 2009<br>Wang *et al.*, 2016<br>Peng *et al.*, 2017 |

Intact gene counts are outside parentheses and pseudogene counts are inside parentheses. <sup>a</sup>pseudogene counts were not addressed explicitly in the study. <sup>b</sup>Zhou *et al.*, 2015 provided counts of truncated models and pseudogenes for ORs and GRs, however these sequences were not published and therefore were not used in building phylogenies.

### *D. alloeum* contains canonical genes involved in reproduction and sex determination

Hymenoptera is an insect order characterized by haplodiploid sex determination, providing an opportunity for studying the evolution of reproductive modes, including transitions from sexual to asexual systems. Meiosis is essential to obligate sexual reproduction, such that loss of sex may be accompanied by the subsequent degradation of meiotic genetic machinery (Schurko & Logsdon, 2008). However, identical sets of meiosis genes in *D. alloeum* (sexual) and *D. muliebre* (asexual) (Tvedte *et al*., 2017) and population genetic data implying that the asexual *D. muliebre* undergoes recombination (Forbes *et al*., 2013) together suggests that asexual wasps retain meiotic production of gametes despite the loss of sexual reproduction. Given the apparent lack of male production in *D. muliebre*, a non-canonical form of meiosis could facilitate the maintenance of genetic variation and promote the persistence of this asexual lineage.

In many hymenopterans, development into male *vs.* female forms is based on allelic states at a single locus, a mechanism known as complementary sex determination (CSD) (van Wilgenburg *et al*., 2006). In *A. mellifera* specifically, sex determination depends on the *csd* gene (Hasselmann *et al*., 2008). We found no evidence of the *csd* locus in *D. alloeum*, however our inability to consistently rear wasps in the laboratory at the current time precludes our ability to definitively rule out CSD as a sex determination mechanism. In CSD and non-CSD hymenopterans, a well-conserved sex determination regulatory cascade includes *transformer* and *doublesex*, both displaying sex-specific splicing (Geuverink & Beukeboom, 2014). We identified single copies of *transformer* and *doublesex* genes in *D. alloeum* (Genbank TBD).

Sex determination genes may be targets of selection in asexual Hymenoptera. Across insects, male production occurs due to alternative splicing of *transformer* rendering the protein nonfunctional, leading to male-splicing of *doublesex*. Conversely, translation of full-length *transformer* into functional protein mediates the splicing of female-specific *doublesex* isoforms (Verhulst *et al*., 2010). RNA-seq read mapping patterns supported sex-specific *transformer* isoforms in *D. alloeum* (Supplementary Figure S10). In all-female *Diachasma* species, we would expect selection to preserve the full-length *transformer* gene. In doublesex, the female isoform in *D. alloeum* is shorter (Supplementary Figure S11), similar to splicing patterns in other insects (Cho *et al*., 2007; Oliveira *et al*., 2009). The single exon specific to males may be subject to future degradation following sex loss in asexual *Diachasma* species.

Additional genes contributing to sex-specific traits (*e.g.* sperm production, pheromones, pigmentation) may be candidates for degradation in asexual wasps (van der Kooi & Schwander, 2014; Kraaijeveld *et al*., 2016). The high quality of *D. alloeum* assembly provides a suitable framework for future studies of the effects of sexual and asexual reproductive modes on patterns of molecular evolution across the wasp genome.

## Materials and Methods

We isolated genomic DNA from a single haploid male and separately from pooled animals of both sexes collected in Fennville, Michigan, USA. A combination of Illumina paired-end, mate pair, and TruSeq Synthetic Long Read (TSLR) libraries were each sequenced on an Illumina HiSeq2000 separately for the single male and pooled samples. Paired-end and mate pair reads were *de novo* assembled using SOAPdenovo2 v2.04 (Li *et al*., 2010) and added TSLR “reads” using PBJelly v2 (English *et al*., 2012). We removed putative microbial contaminant sequences from the assembly that were identified by both BlobTools (Laetsch & Blaster, 2017) and a separate custom pipeline developed by Wheeler *et al*. (2013) and modified as described in Poynton *et al*. (2018). We separately assembled the mitochondrial genome *de novo* using NOVOplasty v2.6.3 (Dierckxsens *et al*., 2017).

We used ten wasps of each sex to generate two (pooled male and pooled female) paired-end RNASeq libraries, and sequenced read libraries using an Illumina HiSeq2500. We combined read datasets and assembled a transcriptome *de novo* with Trinity (Release 2014-04-13) (http://trinityrnaseq.github.io/) (Grabherr *et al.* 2011; Haas *et al.* 2013). Annotation of the *D. alloeum* genome assembly was performed by the NCBI using their Eukaryotic Genome Annotation Pipeline (https://www.ncbi.nlm.nih.gov/genome/annotation_euk/process/), with experimental support from the RNAseq and transcriptome. Manual annotations were added to a *D. alloeum* project on the i5k workspace (https://apollo.nal.usda.gov/diaall/jbrowse/; Poelchau *et al*., 2014). See Supplementary Material online for additional information on genome sequencing, assembly, and annotation.

## Acknowledgements

We thank the W. M. Keck Center for Comparative and Functional Genomics at the University of Illinois at Urbana-Champaign for genomic and RNA library construction and sequencing; Chris Fields at the High Performance Computing for Biology Center at the Roy J. Carver Biotechnology Center at the University of Illinois at Urbana-Champaign for performing the PBJelly scaffolding with TSLR reads; Austin Ward at the Biology Department at the University of Iowa for generating custom scripts for generating nonredundant protein datasets; Samuel Cummings at the Biology Department at the University of Iowa for contributing to annotation of chemosensory genes; and Monica Poelchau at the United States Department of Agriculture-Agricultural Research Service for generating the official gene set for *D. alloeum*. This work was supported by the United States Department of Agriculture/National Institute of Food and Agriculture (A2008-35302-18819 to H.M.R., 2015-67013-23289 to J.L.F.) and the National Science Foundation (DEB-1145355 to A.A.F; DEB-1638997 to J.L.F; DEB1257053 and IOS1456233 to J.H.W.).

## References

Austin A, Dowton M. 2000. The Hymenoptera: an introduction. In Hymenoptera: Evolution, Biodiversity and Biological Control. eds. Austin A, Dowton M. pp 3–16. Csiro Publishing, Clayton, AU.

Benton R, Vannice KS, Gomez-Diaz C, Vosshall LB. 2009. Variant ionotropic glutamate receptors as chemosensory receptors in *Drosophila*. Cell 136:149–162. doi.org/10.1016/j.cell.2008.12.001

Abrahamson WG, Blair CP. 2007. Sequential radiation through host-race formation: herbivore diversity leads to diversity in natural enemies. In Specialization, speciation, and radiation: The evolutionary biology of herbivorous insects. ed. Tilmon KJ. pp188–200. University of California Press, Berkeley, CA, USA

Branstetter M, et al. 2017. Genomes of the Hymenoptera. Curr Opin Insect Sci 25:65–75. doi.org/10.1016/j.cois.2017.11.008

Burke GR, Walden KK, Whitfield JB, Robertson HM, Strand MR. 2018. Whole genome sequence of the parasitoid wasp *Microplitis demolitor* that harbors an endogenous virus mutualist. G3-Genes Genom Genet 8:2875–2880. https://doi.org/10.1534/g3.118.200308

Bush GL. 1966. The taxonomy, cytology, and evolution of the genus *Rhagoletis* in North America (Diptera, Tephritidae). B Mus Compar Zool 134:431–562.

Bush GL. 1994. Sympatric speciation in animals: new wine in old bottles. Trends Ecol Evol 8: 285–288. doi.org/10.1016/0169-5347(94)90031-0

Cho S, Huang ZY, Zhang, J. 2007. Sex_Jspecific splicing of the honeybee *doublesex* gene reveals 300 million years of evolution at the bottom of the insect sex_Jdetermination pathway. Genetics 177:1733–1741. https://doi.org/10.1534/genetics.107.078980

Clyne PJ, et al. 1999. A novel family of divergent seven-transmembrane proteins: candidate odorant receptors in *Drosophila*. Neuron 22:327–338. doi.org/10.1016/S0896-6273(00)81093-4

Croset V, et al. 2010. Ancient protostome origin of chemosensory ionotropic glutamate receptors and the evolution of insect taste and olfaction. PLOS Genet 6:e1001064. doi.org/10.1371/journal.pgen.1001064

Dambroski HR, et al. 2005. The genetic basis for fruit odor discrimination in *Rhagoletis* flies and its significance for sympatric host shifts. Evolution 59:1953–1964. doi.org/10.1111/j.0014-3820.2005.tb01065.x

Dierckxsens N, Mardulyn P, Smits G. 2016. NOVOPlasty: de novo assembly of organelle genomes from whole genome data. Nucleic Acids Res 45:e18. doi.org/10.1093/nar/gkw955

Edgar RC. 2004. MUSCLE: multiple sequence alignment with high accuracy and high throughput. Nucleic Acids Res 32:1792–1797. doi.org/10.1093/nar/gkh340

Elsik CG, et al. 2014. Finding the missing honey bee genes: lessons learned from a genome upgrade. BMC genomics 15:86. https://doi.org/10.1186/1471-2164-15-86

English AC, et al. 2012. Mind the gap: upgrading genomes with Pacific Biosciences RS long-read sequencing technology. PLOS One 7:e47768. doi.org/10.1371/journal.pone.0047768

Feder JL, Forbes AA. 2010. Sequential speciation and the diversity of parasitic insects. Ecol Entomol 35:67–76. doi.org/10.1111/j.1365-2311.2009.01144.x

Forbes AA, Bagley RK, Beer MA, Hippee AC, Widmayer HA. 2018. Quantifying the unquantifiable: why Hymenoptera—not Coleoptera—is the most speciose animal order. BMC Ecol 18:21. doi.org/10.1186/s12898-018-0176-x

Forbes AA et al. 2017. Revisiting the particular role of host shifts in initiating insect speciation. Evolution 71:1126–1137. doi.org/10.1111/evo.13164

Forbes AA, Feder JL. 2006. Divergent preferences of *Rhagoletis pomonella* host races for olfactory and visual fruit cues. Entomol Exp Appl 119:121–127. doi.org/10.1111/j.1570-7458.2006.00398.x

Forbes AA, Powell THQ, Stelinski LL, Smith JJ, Feder JL. 2009. Sequential sympatric speciation across trophic levels. Science 323:776–779. doi.org/10.1126/science.1166981

Forbes AA, Rice LA, Stewart NB, Yee WL, Neiman M. 2013. Niche differentiation and colonization of a novel environment by an asexual parasitic wasp. J Evolution Biol 26:1330–1340. doi.org/10.1111/jeb.12135

Forêt S, Maleszka R. 2006. Function and evolution of a gene family encoding odorant binding-like proteins in a social insect, the honey bee (*Apis mellifera*). Genome Res 16:1404–1413. doi.org/10.1101/gr.5075706

Forêt S, Wanner KW, Maleszka R. 2007. Chemosensory proteins in the honey bee: Insights from the annotated genome, comparative analyses and expressional profiling. Insect Biochem Molec 37:19–28. doi.org/10.1016/j.ibmb.2006.09.009

Geuverink E, Beukeboom LW. 2014. Phylogenetic distribution and evolutionary dynamics of the sex determination genes doublesex and transformer in insects. Sex Dev 8:38–49. doi.org/10.1159/000357056

Grabherr MG, et al. 2011. Full-length transcriptome assembly from RNA-Seq data without a reference genome. Nat Biotechnol 29:644–652. doi.org/10.1038/nbt.1883

Haas BJ, et al. 2013. De novo transcript sequence reconstruction from RNA-seq using the Trinity platform for reference generation and analysis. Nature Protoc 8:1494–1512. doi.org/10.1038/nprot.2013.084

Hasselmann M, et al. 2008. Evidence for the evolutionary nascence of a novel sex determination pathway in honeybees. Nature 454:519–522. doi.org/10.1038/nature07052

Hoede C, et al. 2014. PASTEC: An automatic transposable element classification tool. PLoS ONE. 9, e91929. doi.org/10.1371/journal.pone.0091929

Hood GR, et al. 2015. Sequential divergence and the multiplicative origin of community diversity. Proc Natl Acad Sci USA 112:E5980–E5989. doi.org/10.1073/pnas.1424717112

Kraaijeveld K, et al. 2016. Decay of Sexual Trait Genes in an Asexual Parasitoid Wasp. Genome Biol Evol 8:3685–3695. doi.org/10.1093/gbe/evw273

Laetsch DR, Blaxter ML. 2017. BlobTools: Interrogation of genome assemblies. F1000Research 6:1287. doi.org/10.12688/f1000research.12232.1

Larter NK, Sun JS, Carlson JR. 2016. Organization and function of *Drosophila* odorant binding proteins. Elife 5:e20242.

LaSalle J, Gauld ID. 1993. Hymenoptera: their diversity, and their impact on the diversity of other organisms. In Hymenoptera and Biodiversity. eds. LaSalle J, Gauld ID. pp 1–26. CAB International, Wallingford, UK.

Luo R, et al. 2012. SOAPdenovo 2: an empirically improved memory-efficient short-read *de novo* assembler. Gigascience 1:18. doi.org/10.1186/2047-217X-1-18

Ma WJ, Pannebakker BA, Beukeboom LW, Schwander T, Van de Zande L. 2014. Genetics of decayed sexual traits in a parasitoid wasp with endosymbiont-induced asexuality. Heredity 113:424–431. doi.org/10.1038/hdy.2014.43

Nosil, P. 2012. Ecological Speciation Oxford University Press, New York, NY, USA.

Oliveira DCSG, et al. 2009. Identification and characterization of the *doublesex* gene of *Nasonia*. Insect Mol Biol 18:315–324. doi.org/10.1111/j.1365-2583.2009.00874.x

Pelosi P, Iovinella I, Felicioli A, Dani FR. 2014. Soluble proteins of chemical communication: an overview across arthropods. Front Physiol 5:320. doi.org/10.3389/fphys.2014.00320

Peng Y, et al. 2017. Identification of odorant binding proteins and chemosensory proteins in *Microplitis mediator* as well as functional characterization of chemosensory protein 3. PLOS One 12:e0180775. doi.org/10.1371/journal.pone.0180775

Poelchau M, et al. 2014. The i5k Workspace@ NAL—enabling genomic data access, visualization and curation of arthropod genomes. Nucleic Acids Res 43:D714–D719. https://doi.org/10.1093/nar/gku983

Poynton HC, et al. 2018. The toxicogenome of *Hyalella azteca*: a model for sediment ecotoxicology and evolutionary toxicology. Environ Sci Tech 52:6009–6022. doi.org/10.1021/acs.est.8b00837

Robertson HM, et al. 2018. Genome sequence of the wheat stem sawfly, *Cephus cinctus*, representing an early-branching lineage of the Hymenoptera, illuminates evolution of hymenopteran chemoreceptors. Genome Biol Evol accepted. https://doi.org/10.1093/gbe/evy232

Robertson HM, Gadau J, Wanner KW. 2010. The insect chemoreceptor superfamily of the parasitoid jewel wasp *Nasonia vitripennis*. Insect Mol Biol 19:121–136. doi.org/10.1111/j.1365-2583.2009.00979.x

Robertson HM, Wanner KW. 2006. The chemoreceptor superfamily in the honey bee, *Apis mellifera*: expansion of the odorant, but not gustatory, receptor family. Genome Res 16:1395–1403. doi.org/10.1101/gr.5057506

Schurko AM, Logsdon JM Jr. 2008. Using a meiosis detection toolkit to investigate ancient asexual “scandals” and the evolution of sex. Bioessays, 6:579–589. doi.org/10.1002/bies.20764

Simão FA, Waterhouse RM, Ioannidis P, Kriventseva EV, Zdobnov EM. 2015. BUSCO: assessing genome assembly and annotation completeness with single-copy orthologs. Bioinformatics 31:3210–3212. doi.org/10.1093/bioinformatics/btv351

Stireman JO, Nason JD, Heard SB, Seehawer JM. 2006. Cascading host-associated genetic differentiation in parasitoids of phytophagous insects. P Roy Soc Lon B Bio 273:523–530. doi.org/10.1098/rspb.2005.3363

Tvedte ES, Forbes AA, Logsdon Jr JM. 2017. Retention of core meiotic genes across diverse Hymenoptera. J Hered 108:791–806. doi.org/10.1093/jhered/esx062

van der Kooi CJ, Matthey-Doret C, Schwander T. 2017. Evolution and comparative ecology of parthenogenesis in haplodiploid arthropods. Evolution Let 1:304–316. doi.org/10.1002/evl3.30

van der Kooi CJ, Schwander T. 2014. On the fate of sexual traits under asexuality. Biol Rev 89:805–819. doi.org/10.1111/brv.12078

van Wilgenburg E, Driessen G, Beukeboom LW. 2006. Single locus complementary sex determination in Hymenoptera: an” unintelligent” design? Front Zool 3:1. doi.org/10.1186/1742-9994-3-1

Verhulst EC, van de Zande L, Beukeboom LW. 2010. Insect sex determination: it all evolves around *transformer*. Curr Opin Genet Dev 20:376–383. doi.org/10.1016/j.gde.2010.05.001

Vieira FG, et al. 2012. Unique features of odorant-binding proteins of the parasitoid wasp *Nasonia vitripennis* revealed by genome annotation and comparative analyses. PLOS One 7:e43034. doi.org/10.1371/journal.pone.0043034

Vieira FG, Rozas J. 2011. Comparative genomics of the odorant-binding and chemosensory protein gene families across the Arthropoda: origin and evolutionary history of the chemosensory system. Genome Biol Evol 3:476–490. doi.org/10.1093/gbe/evr033

Walsh BD. 1867. The apple-worm and the apple-maggot. J Hort 2:338–343.

Wang S, et al. 2016. Cloning and expression profile of ionotropic receptors in the parasitoid wasp *Microplitis mediator* (Hymenoptera: Braconidae). J Insect Physiol 90:27–35. doi.org/10.1016/j.jinsphys.2016.05.002

Wang Y, Coleman-Derr D, Chen G, Gu YQ. 2015. OrthoVenn: a web server for genome wide comparison and annotation of orthologous clusters across multiple species. Nucleic Acids Res 43:W78–W84. doi.org/10.1093/nar/gkv487

Weinstock GM, et al. 2006. Insights into social insects from the genome of the honeybee *Apis mellifera*. Nature 443: 931–949. doi.org/10.1038/nature05260

Werren JH, et al. 2010. Functional and evolutionary insights from the genomes of three parasitoid *Nasonia* species. Science 327:343–348. doi.org/10.1126/science.1178028

Wharton RA, Marsh PM. 1978. New world Opiinae (Hymenoptera: Braconidae) parasitic on Tephritidae (Diptera). J Wash Acad Sci:147–167.

Wheeler D, Redding AJ, Werren JH. 2013. Characterization of an ancient lepidopteran lateral gene transfer. PLOS One 8:e59262. https://doi.org/10.1371/journal.pone.0059262

Whitfield JB. 2003. Phylogenetic insights into the evolution of parasitism in Hymenoptera. Adv Parasitol 54:69–100.

Zhang S, Zhang Y, Su H, Gao X, Guo Y. 2009. Identification and expression pattern of putative odorant-binding proteins and chemosensory proteins in antennae of the *Microplitis mediator* (Hymenoptera: Braconidae). Chem Senses 34:503–512. doi.org/10.1093/chemse/bjp027

Zhou X, et al. 2015. Chemoreceptor evolution in hymenoptera and its implications for the evolution of eusociality. Genome Biol Evol 7:2407–2416. doi.org/10.1093/gbe/evv149

