## Supplementary Material online for "Genome of the parasitoid wasp *Diachasma alloeum*, an emerging model for ecological speciation and transitions to asexual reproduction"

**Supplemental Material**

### Experimental Procedures

#### Biological Material

We collected fallen fruit from five downy hawthorn trees (*Crataegus mollis*) historically highly infested with both *R. pomonella* (Diptera: Tephritidae) and *D. alloeum* (Hymenoptera: Braconidae) in Fennville, MI (42.597307, -86.151498) in August 2009 and 2010. Infested fruit were transported to the greenhouse at the University of Notre Dame, and placed in wire cages above black planter trays that caught late-instar fly larvae as they exited fruits. As fly larvae pupated, we moved them into Petri dishes containing moist vermiculite, held them at room temperature (21 ± 2ºC) for 10 days, and then moved them into a 4°C refrigerator for 4 months to simulate overwintering conditions (Forbes *et al*., 2009; Hood *et al*., 2015). After overwinter, we removed Petri dishes from the cold, placed the dishes in several 15 cm x 15cm x 25 cm Plexiglas cages supplied with water and a diet of honey mixed with brewer’s yeast inside a rearing chamber held at 24ºC with a 14:10 light:day cycle. Cages were monitored daily for eclosing *R. pomonella* flies and wasps. Upon eclosion, adult wasps were kept alive for 3-7 days, and then frozen at -80°C. At a later date, preserved wasps were sexed and identified to the species level using the key of Wharton & Yoder (2015).

#### DNA isolation, library preparation, sequencing, and genome assembly

We selected a single male wasp and pooled female animals for DNA sequencing. Briefly, we homogenized whole adult animals in liquid nitrogen and lysed tissue in a SDS solution overnight with Proteinase K. We treated the homogenate with RNaseA, and we collected proteins/debris after high-salt precipitation and centrifugation. After ethanol precipitation, we resuspended the DNA in 10 mM Tris and subsequently evaluated extracted DNA on an agarose gel and by Qubit quantification.

We generated the following libraries for sequencing: a 500 bp shotgun library from a single male wasp, a 1.5 kbp shotgun library from four pooled female wasps, a 5 kbp mate-pair library using a four female + one male pooled sample, and 10 and 20 kb insert mate-pair libraries using DNA from 50 pooled mixed sex wasps. Pooled samples were used to achieve the minimum DNA mass needed for library preparation. We prepared the 500 bp and 1.5 kb shotgun libraries with Illumina TruSeq DNAseq Sample Prep kits. We prepared the 5 and 10 kb mate-pair libraries using a similar protocol, however we used a custom linker to ligate between the fragment ends to facilitate mate-pair recovery. We constructed the 20 kb mate-pair library with an Illumina Nextera Mate-Pairs Sample Prep kit. We sequenced all libraries for 100 cycles on an Illumina HiSeq2000 using Illumina TruSeq SBS Sequencing kit v3. Bases were called with Casava v1.8 and the reads are available from the Short Read Archive at NCBI (SRR2042503, SRR2046752, SRR2042775, SRR2043489, SRR2043491, SRR2043616, SRR2043618, SRR2043726, SRR2041626).

We filtered mate-pair libraries for properly-oriented reads of the appropriate insert size and uniqueness using in-house custom pipeline scripts. We trimmed raw Illumina reads on 5' and 3' ends nucleotide-bias and low-quality bases using the FASTX Toolkit (http://hannonlab.cshl.edu/fastx_tookit/). Next, we error-corrected trimmed reads by library with Quake (Kelley *et al*., 2010), counting 19-mers. To minimize haplotype reconstruction issues during the initial *de novo* assembly, we used SOAPdenovo v2.04 (Luo *et al*., 2012) with K=49 to assemble the 500 bp-insert shotgun library reads sequenced from an individual haploid male. Following this step, we used SOAPdenovo to perform scaffolding with iteratively longer-insert shotgun and mate-pair libraries and use of GapCloser v1.12 to close gaps generated in the scaffolding (Luo *et al*., 2012).

In addition, to improve scaffolding of the genome assembly, we generated an Illumina TruSeq Synthetic Long Read (TSLR) library with a TSLR Sample Prep kit from the same pooled DNA sample. We sequenced the library for 100 cycles on a HiSeq2000 and the resultant read data was analyzed by Illumina using their TruSeq Long Read Assembly Application in “BaseSpace”. We added the resultant TSLR “reads” to the assembly using PBJelly v2 (English *et al*., 2012).

#### RNA isolation, RNAseq library preparation, sequencing, and transcriptome assembly

We prepared separate homogenized samples for ten wasps of each sex in 1ml Trizol in glass tissue grinders and filtered tissue over a Qiagen Qiashredder column. We extracted the homogenates with chloroform and the precipitated RNA with linear polyacrylamide (10mg/mL) and isopropanol. Next, we washed RNA pellets with 75% ethanol and resuspended in RNase-free water. We quantified RNA with a Qubit RNA Broad Range Assay Kit on a Qubit fluorometer (Life Technologies). We visualized RNA using ethidium bromide on a 1.0% agarose gel. We then prepared RNAseq libraries from an average cDNA fragment size of 250 bases using the Illumina TruSeq Stranded RNAseq Sample Prep kits. We individually barcoded the two libraries and quantitated libraries using qPCR before pooling and sequencing from both ends with TruSeq SBS Sequencing kit v4 for 100 cycles on a HiSeq2500 instrument. Bases were called with Casava v1.82. We trimmed reads as for DNA sequencing and assembled the combined trimmed reads from males and females with Trinity (Release 2014-04-13) (http://trinityrnaseq.github.io/) (Grebherr *et al*., 2011; Haas *et al.*, 2013). The raw RNAseq reads were submitted to the Sequence Read Archive (SRA) at NCBI (accession numbers SRR2041626 and SRR2040481), while the Trinity transcriptome assemblies were submitted to the Transcriptome Shotgun Assembly database (accession numbers GECN00000000.1).

#### Contaminant removal and lateral gene transfer analysis

Identification of contaminating sequences in a genome assembly remains a challenging task, and we used multiple strategies to remove contaminants from the *D. alloeum* genome. First, our submission of the *de novo* assembled *D. alloeum* genome included a screen for vector, bacterial, and library adaptor sequences (<https://www.ncbi.nlm.nih.gov/tools/vecscreen/contam/>) prior to the submission of the assembly to the Eukaryotic Genome Annotation pipeline at the NCBI (<https://www.ncbi.nlm.nih.gov/genome/annotation_euk/process/>). We removed these contaminating sequences from the assembly prior to acceptance by NCBI.

Second, we generated a blobplot using BlobTools v1.0 (DOI 10.5281/zenodo.845347, Laetsch & Blaxter, 2017). Using the ‘taxify’ module, we conducted BLASTn comparisons between the *D. alloeum* scaffolds in the assembly and the NCBI nt database, generating a ‘hits’ file. We used bwa-mem (Li *et al*., 2013) to align paired-end reads from the 500 bp insert library to the *D. alloeum* genome to generate a BAM coverage file. We submitted the assembly file, ‘hits’ file, and BAM coverage file using the ‘create’ module to produce a database output file, which we then visualized using the ‘view’ module. We further inspected scaffolds with hits to non-Arthropod identifiers, and we identified sequences with aberrant GC content and coverage as putative contaminants.

Third, we employed a custom pipeline developed by Wheeler *et al*. (2013) and further refined as described in Poynton *et al*. (2018) to identify contaminants and potential lateral gene transfer (LGT) events. For each scaffold in the *D. alloeum* assembly, we characterized the number and size of bacterial hits, bacterial breadth of coverage, scaffold size, and the genus of the best bacterial hit. The pipeline uses scaffold size and bacterial coverage to assign scaffolds as putative contaminants.

As a conservative measure, we removed scaffolds that were assigned as putative bacterial contaminants by either BlobTools or the custom Wheeler and Werren pipeline. Manual inspection of discrepant assignments revealed one scaffold of interest (RefSeq accession NW_015145431.1), which was called as bacterial by BlobTools but not by the custom pipeline, and additional investigation revealed a potential LGT event (see Results & Discussion). Our curation of this scaffold followed the methods described in Poynton *et al*. (2018). This scaffold was split into 1kb fragments, followed by assignment of each fragment to eukaryotic and bacterial bitscores based on comparisons with eukaryotic gene models and bacterial genome sequences, respectively. We identified potential LGT events as regions with high bacterial bitscores and low eukaroytic bitscores flanked by regions with the opposite pattern, *i.e.* high eukaryotic bitscores and low bacterial bitscores. To further characterize genes and in this scaffold, we used regions of interest as queries for BLASTn and BLASTx searches against nt/nr databases with the “bacteria” organism filter. We also submitted this sequence to CENSOR (Kohaney *et al*., 2006) to assess whether transposable elements have accumulated near LGT regions.

#### Characterization of ultraconserved elements using BUSCO

The BUSCO tool (Simão *et al*., 2015) estimates the completeness of a genome assembly by providing a quantitative measure of the presence of single-copy orthologs. We retrieved Arthropod and Hymenoptera ortholog datasets from OrthoDBv9 (Zdobnov *et al*., 2017) which contain genes present in > 90% of the species within those respective groups in OrthoDB. Using default parameters, we ran BUSCO v3 in genome mode on the *D. alloeum* assembly. We downloaded genome assemblies for other hymenopteran species from NCBI (*A. mellifera*, *N. vitripennis*, *M. demolitor*, Table 2). Using the same run parameters, we identified the presence of arthropod and hymenopteran BUSCOs for published hymenopteran genome assemblies for comparative purposes.

#### Transposable element analysis

We ran RepeatModeler (Smit *et al*., 2015, v1.0.8), utilizing RECON (Bao & Eddy, 2002) and RepeatScout (Price *et al*., 2005), on the *D. alloeum* genome assembly to *de novo* assemble repetitive sequence families. We used the accompanying RepeatModeler program, RepatClassifier to assign class and family classifications to these repeats based on similarity to RepBase (Bao *et al*., 2015, v22.10). To more thoroughly classify transposable elements (TEs) in the RepeatModeler output we used PASTEClassifier (Hoede *et al*., 2014, v1.0) to classify TE sequences in the RepeatModeler output based on structural features such as terminal inverted repeats (TIRs), long terminal repeats (LTRs), polyA tails and coding regions such as transposase and reverse transcriptase. Additionally, we used the options for similarity comparison to TEs in RepBase and the Pfam hidden Markov model protein domain profiles (Finn *et al*., 2016, Pfam27.0_GypsyDB). We combined the results from these programs to reclassify sequences with discrepancies between the RepeatClassifier results and the multiple levels of evidence from PASTEClassifier for final classifications according to hierarchical scheme outlined by Wicker *et al*. (2007) where TEs belong to either DNA transposon or retrotransposon class, an order within that class (e.g. long terminal repeat, LTR, retrotransposon) and superfamily (*e.g.* LTR *Gypsy*) where possible. We performed *de novo* repeat detection and abundance estimation for *Nasonia vitripennis* (GCA_000002325.2_Nvit_2.1) and *Apis mellifera* (GCA_000002195.1_Amel_4.5) for comparative purposes.

We used RepeatMasker (Smit *et al*., 2010, version 4.0.7) to estimate the abundance of TEs in the *D. alloeum*, *N. vitripennis*, and *A. mellifera* genome assemblies and the utility script buildSummary.pl to summarize these results by TE superfamily, order, and class. We also used parseRM.pl (<https://github.com/4ureliek/Parsing-RepeatMasker-Outputs>) to extract and bin divergence data from the RepeatMasker output by TE class and then generated repeat landscapes to visualize past TE activity.

#### Manual gene annotation

*Oxidative phosphorylation genes*

We used NOVOplasty v2.6.3 (Dierckxsens *et al*., 2017) to assemble the mitochondrial genome of *D. alloeum*. To *de novo* assemble the mitochondrial sequence, we supplied raw reads from the 500bp insert library using the cytochrome C oxidase 1 gene (cox1) from a different individual *D. alloeum* wasp (Forbes *et al*., 2009) (Genbank EU881670.1) as a seed input. We used MUSCLE (Edgar, 2004) to align mitochondrial sequence regions of interest with *Diachasmimorpha longicaudata* sequences (Wei *et al*., 2010). Using MUSCLE alignments and *de novo* predictions of protein coding genes, rRNAs, and tRNAs from the MITOS web server (Bernt *et al*., 2013), we annotated 13 protein-coding genes, two rRNA sequences, and 22 tRNAs.

We searched for 68 nuclear OXPHOS genes in the *D. alloeum* assembly using queries from *N. vitripennis*, *A. mellifera*, and *D. melanogaster* (Porcelli *et al*., 2007; Gibson *et al*., 2010). To confirm putative nuclear OXPHOS genes, we performed a reciprocal BLAST search using annotated proteins from insects as queries to search *D. alloeum* genome using tBLASTn, and subsequently performed BLASTx against the NCBI nr database to confirm the identity of candidate *D. alloeum* OXPHOS sequences. To assist with annotations, we incorporated predictions from GeneWise (Birney *et al*., 2004) using orthologous OXPHOS proteins from other insects. We then assigned genes to the OXPHOS enzyme complexes I-V based on information from MitoComp2 (<http://www.mitocomp.uniba.it/>; Porcelli *et al*., 2007).

*Chemosensory genes*

To characterize chemosensory gene families in D. alloeum, we used the reciprocal BLAST strategy described above. We obtained protein sequences from five major families (odorant receptors – ORs; gustatory receptors – GRs; ionotropic receptors – IRs, odorant binding proteins – OBPs; chemosensory proteins – CSPs) from published data from *N. vitripennis*, *A. mellifera*, *M. demolitor*, and *M. mediator*. Table 3 provides source information for chemosensory gene sequences. We conducted tBLASTn searches of the *D. alloeum* genome using hymenopteran proteins as queries. For gene families with highly divergent sequence, *i.e.* ORs, GRs, IRs, we used the BLOSUM45 substitution matrix to execute searches. All hits with an e-value less than 1e-5 were manually inspected. We performed BLASTx searches on putative *D. alloeum* chemosensory gene nucleotide sequences against the NCBI nr database and retained genes with top hits matched to the appropriate gene family.

To construct gene models, we used the annotation features of the *D. alloeum* WebApollo portal (<https://apollo.nal.usda.gov/diaall/jbrowse/>). We compared hymenopteran proteins with the best tBLASTn score to *D. alloeum* nucleotide sequences using GeneWise (Birney *et al*., 2004). Protein sequences were aligned using the CLUSTALW plugin on the Geneious bioinformatics software platform (Kearse *et al*., 2012). We retained sequences with at least ~50% of the characteristic length of the protein family (ORs/GRs/IRs > 200 aas, OBPs/CSPs > 75 aas), and performed iterative rounds of alignment and reannotation to fix poorly aligning regions. Final gene models included start sites, stops sites, and exon-intron boundaries. Functional classification was assigned to gene models using InterProScan (Zdobnov & Apweiler, 2001).

After obtaining final models for chemosensory genes, we constructed protein alignments using CLUSTALW followed by trimming of poorly aligning regions using trimAl v1.2 running in “strict” (IRs) or “gappyout” (ORs, GRs, OBPs, CSPs) modes (Capella-Gutiérrez *et al*., 2009). We determined the best-fit substitution model for each family using the IQ-TREE webserver ([http://iqtree.cibiv.univie.ac.at](http://iqtree.cibiv.univie.ac.at/); Trifinopoulos *et al*., 2016) which served as the substitution matrix for maximum likelihood phylogenies, produced here using RAxML (Stamatakis, 2006). To evaluate node support, we conducted 1000 bootstraps in RAxML runs. Phylogenies and their support values were visualized using FigTree v1.4.3 (<http://tree.bio.ed.ac.uk/software/figtree/>).

We followed the naming convention for ORs and GRs in *D. alloeum* largely based on previously published works on chemoreceptors in model hymenopteran species (Robertson & Wanner, 2006; Robertson *et al*., 2010). Genes were named with a four-letter prefix corresponding to the species name, followed by the gene family name, followed by a number (*e.g.* DallOr1, DallOr2, DallOr3, etc. and DallGr1, DallGr2, DallGr3, etc.). DallOrs and DallGrs were named in ascending order corresponding to sequence orthologs in *N. vitripennis* and *A. mellifera*, however extensive gene duplication and loss in these gene families prevented a consistent naming scheme.

*D. alloeum* IR family names followed the conventions outlined in Croset *et al.* (2010). We categorized and named IRs into “antennal IRs” and “divergent IRs.” The “antennal IRs” represent a collection of well-conserved IRs that were first characterized in the antenna of *D. melanogaster* (Benton *et al*., 2009). We generated alignments for IRs using sequences from *D. melanogaster* and various hymenopteran species (tree not shown). We assigned the same name to unambiguous orthologs of *D. melanogaster* “antennal IRs” (e.g. DmelIr8a = DallIr8a). In the event that multiple copies of an IR sequence ortholog exist for *D. alloeum*, we assigned the genes with the same base name followed by a point and a number (e.g. DallIr64a, DallIr64a.2) We numbered *Diachasma*-specific IRs and those with unclear orthology in ascending order, starting with DallIr101. The “divergent IRs” have low intraspecific and interspecific sequence identity and are expressed in gustatory organs of *D. melanogaster* (Croset *et al*., 2010).

We adapted naming conventions for OBPs and CSPs proposed by Forêt & Maleska (2006) for *A. mellifera* genes. We used a four letter Dall prefix followed by a three letter abbreviation that indicates what family the gene belongs to (OBP/CSP). We appended numbers at the end of each gene, with tandemly-arranged genes receiving sequential numbers.

*Sex determination genes*

To search for orthologs of *doublesex*, *transformer*/*feminizer*, and *csd* in the *D. alloeum* genome, we used the reciprocal BLAST strategy described above for OXPHOS and chemosensory genes. Genes used as initial BLAST queries are provided in Supplementary Table S1.

### Results & Discussion

#### Annotation of putative contaminant sequences in *D. alloeum genome assembly*

The *de novo* assembled *D. alloeum* genome contained 93 scaffolds with high sequence similarity to the α-proteobacterium *Wolbachia*, which is unsurprising given this obligate endosymbiont is prevalent among arthropods (Werren *et al*., 1997; Hilgenboecker *et al*., 2008). Indeed, the genome sequence of the *Wolbachia* endosymbiont of *D. alloeum* was recently assembled using raw read data from this study (Pascar & Chandler, 2018). No other bacterial contaminants were removed from the NCBI screening process.

After removing *Wolbachia* sequences, the draft *D. alloeum* genome assembly contained 3,968 scaffolds (Table 1). Of these, 508 scaffolds were assigned non-Arthropod taxonomic identifiers by BlobTools, including 491 having a Proteobacteria taxonomic identifier. Many of these hits contained higher GC content than a typical Arthropod scaffold, providing additional evidence that these are contaminating sequences (Supplementary Figure S1). The complete output of BlobTools is provided in Supplementary File S1.	The custom pipeline (JH Werren) described in Poynton *et al*. (2018) identified 635 putative bacterial scaffolds in the *D. alloeum* assembly. A combined 656 *D. alloeum* scaffolds were assigned as non-Arthropod by either pipeline, indicating consistent contamination calls across pipelines, and these are included in Supplementary File S1. Among these are a large number of scaffolds from bacteria related to *Pectobacterium*, as well as *Soldalis*-like, *Arsenophonus*-like and *Citrobacter*-like bacteria. These putative contaminants represent sequences from potential set of microbial associates of *D. alloeum* (metagenomic data), although we currently cannot rule out contamination from the host, food, or other sources. Further work is needed to determine their status relevant to *D. alloeum* biology.

While most scaffolds with bacterial matches are small and most likely represent contamination, one exception was a 366 kb scaffold (RefSeq accession NW_015145431.1) assigned as bacterial using BlobTools, but not in the Wheeler and Werren pipeline. Given only ~2% of the scaffold length had bacterial hits, this sequence likely represents a case of LGT. In addition, read depth for the rickettsial-like regions of the scaffold are similar to other genomic scaffolds, consistent with their nuclear origin. BLASTn searches of this region produced hits to genome sequences from the bacterial genus *Rickettsia*, and BLASTx searches had hits to two *Rickettsia* genes: phenylalanine – tRNA ligase subunit beta and pyruvate, phosphate dikinase. Both gene-encoding regions in *D. alloeum* have nonsense mutations, and there was a frameshift mutation in the phenylalanine – tRNA ligase subunit in the wasp scaffold. We also identified numerous matches to eukaryotic transposable elements in this region, including DNA transposons, LTR and non-LTR retrotransposons (Supplementary File S1). The pattern of interspersed bacterial and eukaryotic TEs is a signature of a large LGT that has subsequently begun to accumulate TE insertions, similar to the large Wolbachia LGT detected in the fly *Drosophila ananassae* (Hotopp *et al*., 2007). Taken together, the evidence supports the view that scaffold NW_015145431.1 contains a LGT from *Rickettsia* that has subsequently pseudogenized by TE insertions, point mutations and indels. The LGT event is likely relatively recent, as the regions in the *D. alloeum* assembly share > 90% nucleotide identity with their corresponding *Rickettsia* hits. LGT can be a source of evolutionary novelty, and the movement of DNA from bacteria to diverse eukaryotes has been documented (*e.g.* Hotopp *et al*., 2007; Crisp *et al*., 2015). However, it is unlikely this LGT event actively confers benefits to *D. alloeum*, as these sequences are degraded in the wasp genome. The failure of *D. alloeum* RNASeq data to map to these regions suggests these genes are not transcribed and provides additional evidence for their inactivity in the wasp genome.

#### Characterization of BUSCO genes in *D. alloeum*

We identified 1053 of 1066 (~99%) arthropod BUSCOs and 4162 of 4415 (~94%) of hymenopteran BUSCOs in the *D. alloeum* genome assembly (Supplementary Tables S2, Supplementary Table S3). Details of the BUSCO results are provided in Supplementary File S2.

The recovery of complete BUSCOs is comparable to amounts recovered for *A. mellifera*, *N. vitripennis*, and *M. demolitor*. As the number of genes contained by > 90% of OrthoDB taxa is expected to be higher in a narrower taxonomic group (*e.g.* Hymenoptera relative to Arthropoda), we would also expect spurious gene loss and duplication in species for this expanded gene set. As such, we recovered a higher number of duplicate, fragmented, and missing Hymenoptera BUSCOs relative to Arthropoda BUSCOs. When we combined missing and fragmented BUSCOs into an “incomplete” category, we found a considerable percentage of Arthropoda (3/13, 23%) and Hymenoptera (93/253, 37%) BUSCOs that were incomplete in the *D. alloeum* assembly that were also incomplete in at least one other hymenopteran surveyed. The most frequent pattern we observed was genes that were incomplete in *M. demolitor*, which we identified in 3/13 (23%) Arthropoda BUSCOs and 78/253 (31%) in Hymenoptera BUSCOs that also were incomplete in *D. alloeum*. Although assembly fragmentation may preclude the ability of BUSCO to retrieve the appropriate sequences, an alternate explanation is that these genes were lost prior to the divergence between *Diachasma* and *Microplitis*. Overall, the identification of BUSCOs in the *D. alloeum* genome is comparable and often superior to gene content present in the scaffold assembly of *M. demolitor* and the chromosomal-level assemblies of *A. mellifera* and *N. vitripennis.*

#### Transposable elements content in *D. alloeum*

Transposable elements identified in *D. alloeum* are summarized in Supplementary Table S4, containing 367 TE families and 1048 “Unknown” repeat families. These numerous unclassified sequences may include novel TE families but they lack typical structural and coding sequences of TEs or similarities to already classified TEs. We found that DNA transposons have over 100 more distinct sequence families than retrotransposons, the majority of being miniature inverted repeats (MITEs), which are small non-autonomous cut-and-paste transposons with terminal inverted repeats (TIRs). Despite this discrepancy in diversity of families, DNA and retrotransposons contribute similar amounts of sequence to the *D. alloeum* genome (10% DNA, 8% retro). The most abundant superfamily among TIR DNA transposons is TcMar (1% of the genome), while unclassified TIR transposons constitute 4% of the genome. LTR retrotransposons are more abundant than LINEs (5% *vs*. 2% of the genome) and SINEs are the least abundant group of retrotransposons (0.03% of the genome). Interestingly, *Mavericks* (*i.e.* *Polintons*) are the most abundant superfamily (3% of the genome), and the followed closely by the LTR *Gypsy* group (3% of the genome). *Mavericks* are notable for being exceptionally large (up to 20 kb in length), likely related to DNA viruses, and moving via replicative transposition without an RNA intermediate (Wicker *et al*., 2007).

TE activity has been ongoing and sustained for all of the main TE classes though *Helitrons* and *Penelope* retrotransposons appear to have become inactive recently in *D. alloeum*, based on their absence from the lowest divergence bins (Supplementary Figure S2). We also noticed that among the LINEs, *R1* was previously the most abundant until its apparent inactivation with *LOA* activity following and becoming the sole detectable active LINE (Supplementary Figure S3).

We found that *D. alloeum* has markedly more repetitive DNA in its genome than *Nasonia* (Supplementary Table S5), though the largest discrepancy is in “Unknown” sequences. Though the TE portion of the *D. alloeum* is nearly twice that of *N. vitripennis* genome, the two species have a similar diversity of TE families in terms of the superfamilies present and number of distinct TE families (367 in *D. alloeum* vs. 338 in *N. vitripennis*) though the *N. vitripennis* genome is more populated by retrotransposons than DNA transposons and its LINEs and LTRs constitute equal genomic portions*.* The inferred TE activity in *N. vitripennis* is broadly similar to that of *D. alloeum* in that the main TE orders have all had activity throughout much of the genome’s evolutionary history (Supplementary Figure S4). In contrast to the similarities between TEs in these two wasps, the *A. mellifera* genome is almost devoid of TEs, a hallmark characteristic of honeybee genomes (Park *et al*., 2015). Not only do TEs make up a considerably small portion of the *A. mellifera* genome (0.37% of the genome) but TE diversity is also low, as indicated by the relatively few families and reduced contribution from retrotransposons, represented by only a single *Copia* LTR family (Supplementary Table S6). Unlike *D. alloeum* and *N. vitripennis* TIR DNA transposons are by far the most abundant type of TE in the *A. mellifera* genome, with TcMar accounting for nearly all of its classified TEs content (13/18 families and 95% of TE sequence). The *A. mellifera* genome does, however, have the largest genomic proportion of simple repeats among these three species at 4.34% of the genome.

#### Manual gene annotation in *D. alloeum*

*Oxidative phosphorylation genes*

We retrieved one large contig of 15,936 bp containing a complete set of 13 protein-coding genes, 22 tRNAs, and two RNAs from the NOVOPlasty output, representing the mitochondrial genome of *D. alloeum* (Supplementary File S3). The total sequence length falls within the range of other complete braconid wasp genomes (15,000-19,000 bp; Wei *et al*., 2010). We discarded two small contigs < 2,000 bp corresponding to mitochondrial locations surrounding the A+T-rich region. By default, NOVOPlasty lists possible combinations of small contigs to produce a complete circular genome. Without evidence *a priori* to favor certain arrangements over others, we chose to discard these small contigs and retain only the single high-confidence contig.

All *D. alloeum* mitochondrial protein-coding genes and RNAs in have expected sizes and are complete. *D. alloeum* sequences have an identical spatial arrangement to the braconid wasp *Diachasmimorpha longicaudata* (Supplementary Figure S7; Wei *et al*., 2010). All tRNAs had typical cloverleaf structures predicted using the MITOS web server (<http://mitos.bioinf.uni-leipzig.de/index.py>; Bernt *et al*., 2013). No secondary structure analyses were performed on rRNAs, but BLASTn searches of these sequences confirmed their identity.

We annotated 65 of 68 nuclear-encoded mitochondrial genes in the *D. alloeum* genome (Supplementary File S3). The three missing genes (B15 and MNLL in complex I, polypeptide VIC in complex IV) are all present in *A. mellifera*, but not found in the *N. vitripennis* genome (Porcelli *et al*., 2007; Gibson *et al*., 2010) indicating these genes may have been lost after the emergence of parasitoidism in Hymenoptera. Of these 65 genes, we characterized 62 genes as full-length: amino acid sequences are intact and have similar lengths to homologs in *N. vitripennis*, *A. mellifera,* and *D. melanogaster*. We generated truncated models for the remaining three genes. We also found gene duplicates for four genes (flavoprotein and iron-sulfur subunits in complex II, subunit IV and polypeptide VIA in complex IV). These duplicates have RNASeq support and considerable homology (>30% sequence identity) with duplicated sequences in *A. mellifera* and *D. melanogaster.*

*Diachasma alloeum* is a potentially powerful natural system to evaluate patterns of molecular evolutionary rate in mitochondrial and nuclear genes in the OXPHOS pathway. Recently, it has been demonstrated that both mitochondrial and nuclear OXPHOS genes display higher rates of amino acid substitutions in Hymenoptera relative to other insect orders (Li *et al*., 2017). Accelerated molecular evolution in nuclear OXPHOS genes may be a consequence of positive selection acting on compensatory changes in response to mutation accumulation in mitochondrial OXPHOS genes and/or reflect relaxation of functional constraint on certain regions of these proteins (Rand *et al*., 2004; Zhang & Broughton, 2013). Elevated rates in some nuclear-encoded OXPHOS genes have been implicated in mitochondrial-nuclear incompatability between closely-related species of *Nasonia* (Gibson *et al*., 2010). The OXPHOS gene set could be used to assess the presence/absence of postzygotic reproductive barriers between populations of *D. alloeum* utilizing different host plants (Forbes *et al*., 2009). In addition, co-transmission of nuclear and mitochondrial genomes as a single unit in asexuals may influence patterns of mutation accumulation in both genomes, and comparative studies have demonstrated higher mutational loads in asexual mitochondrial genomes relative to their sexual counterparts (Normark & Moran, 2000; Neiman *et al*., 2009; Sharbrough *et al*., 2018). In another species of *Diachasma* that has experienced recent loss of sex, various phenotypically different lineages of asexual wasps based on distinct mtDNA haplotypes have been identified (Forbes *et al*., 2013). The extent to which genetic variation is generated and transmitted in asexual mitochondrial genomes may provide insight into the adaptability and long-term persistence of this asexual species.

*Chemosensory genes*

We annotated a total of 321 putative chemosensory genes in *D. alloeum*. All models had confirmed BLASTx hits with members of their respective protein families, and most intact (non-pseudogene) models had at least one characteristic InterPro domain associated with the protein family with which it belongs. Detailed information for chemosensory genes is available in Supplementary File S4.

We found 201 odorant receptors in the *D. alloeum* genome (Table 3). The number of *D. alloeum* ORs is comparable to *A. mellifera* (174 AmelOrs), *N. vitripennis* (301 NvitOrs), and *M. demolitor* (222 MdemOrs), consistent with the expansion of this gene family in Hymenoptera (Robertson & Wanner, 2006; Robertson *et al*., 2010; Zhou *et al*., 2015). Among *D. alloeum* ORs, 187 were intact, while 14 are putative pseudogenes. The 187 intact ORs may be an overestimate, as this includes some truncated models with missing N-terminal or C-terminal ends that may actually represent pseudogenes. Since we did not include any fragments less than half of the length of a typical OR gene (< 200 amino acids), it is unlikely that truncated annotations represent the N and C termini of the same gene. Of the intact genes annotated in this study, 69 models were correctly predicted by the NCBI pipeline, 71 were changed, and 47 were unannotated. All intact ORs had InterPro domains characteristic for the OR family (IPR004117). The majority of DallOrs (159/201) are present in tandem repeats, an observation consistent with spatial patterns of these genes in other hymenopterans (Robertson & Wanner, 2006; Robertson *et al*., 2010; Zhou *et al*., 2015).

The phylogeny of OR proteins is shown in Supplementary Figure S6. The sole 1:1 ortholog group contained DallOr1, MdemOr1, NvOr1, and AmOr2; these orthologs of DmOr83 serve as OR coreceptors (ORCOs) and are widely conserved across insects (Krieger *et al*., 2003). All other DallOrs exhibit complicated patterns of gene duplication and loss relative to ORs in other surveyed hymenopterans. We found OR subfamilies that were present in the three parasitoid wasp species (*D. alloeum*, *M. demolitor*, *N. vitripennis*) that were either absent or greatly reduced in *A. mellifera* (Supplementary Figure S6A). Conversely, the tandem duplication in AmelOr1-61 represents a substantial expansion relative to the orthologous 15 gene tandem repeat in *D. alloeum* (DallOr2-16, Supplementary Figure S6B). The relationships among members in this tandemly repeated subfamily are complicated by the lack of monophyly among species-specific ORs, and orthologous sequences in *N. vitripennis* are located on separate chromosomal locations (Robertson *et al*., 2010). In comparisons of parasitoid wasps, there are orthologous group gene ratios of one-to-many ORs, such as the DallOr196 placement with MdemOr155-167 (Supplementary Figure S6C). There are also major subfamily expansions completely absent in *D. alloeum*, such as the 75-gene group of *N. vitripennis* (Supplementary Figure S6D).

The OrthoVenn analysis provided details on gene clusters putatively specific to *D. alloeum*. We re-analyzed these clusters after manual annotation of *D. alloeum* ORs. We found one cluster (DallOr114-122) that had no obvious orthologous subfamily in other species in the phylogenetic tree and may represent a *bona fide* expansion in a recent wasp lineage (Supplementary Figure S7E). However, we also identified multiple examples of clusters that grouped with subfamilies from other species, *e.g.* DallOr192-197 and DallOr17-26 (Supplementary Figure S6F, Supplementary Figure S6G). Identification of synteny on one side of these tandem duplicates between *D. alloeum* and other hymenopterans provides strong support that these are orthologous gene clusters. Overall, it is possible that *D. alloeum*-specific cluster identification and associated GO-term enrichment is influenced by large sequence divergence between hymenopteran OR sequences. Assessing legitimate expansions in *D. alloeum* is complicated by the indeterminate orthology of clusters and fragmentation of genome assemblies. Nevertheless, the expansion of OR subfamilies consistent with other parasitic wasps reflects the complex chemical ecology underlying host finding and mating behaviors in *D. alloeum.*

The gustatory receptor (GR) family composition varies considerably across Hymenoptera, and the 40 GRs we identified in *D. alloeum* is larger than the 10 GRs identified in *A. mellifera* (Robertson & Wanner, 2006) but smaller than the 86 identified in *M. demolitor* and 58 identified in *N. vitripennis* (Table 3; Robertson *et al*., 2010, Zhou *et al*., 2015). DallGr28 was the only gene identified as a putative pseudogene, and among the 39 intact genes, only four were accurately predicted by NCBI. We changed seven models and annotated 28 new models in this study. All annotated GRs possess the IPR013604 InterPro domain specific to seven-transmembrane chemoreceptors. A majority (23 of 40) of these genes are arranged in tandem repeats in *D. alloeum*, consistent with observations of spatial patterns of GRs in *N. vitripennis* (Robertson *et al*., 2010). In addition, while not quantified, there was a notable representation of DallGrs not assigned to tandem groups that were near edges (> 50,000 bp) or on scaffolds < 10 kbp.

A phylogeny of DallGrs, MdemGrs, NvGrs, and AmGrs is shown in Supplementary Figure S7. Simple 1-to-1 orthologous groupings were recovered for DallGr1 and DallGr2, which are putative sugar receptors, and DallGr3, which is related to the fructose receptor of flies (Robertson *et al*., 2010). We also recovered DallGr5 as a simple ortholog with the three other hymenopterans (AmelGr6, NvitGr6, MdemGr76). DallGr4 has high sequence identity (> 70%) with a recently duplicated GR in *N. vitripennis* (NvitGr4-5), although surprisingly this group did not include an ortholog from *M. demolitor*. DallGr6 has strong support for placement with tandem duplicates in *A. mellifera* (AmelGr8-9) and *N. vitripennis* (NvitGr8-9), although the orthology of these genes is unclear.

Similar to observations in ORs, extensive gene duplication and loss is present in the GR phylogeny. There is strong support for a subfamily expansion containing three truncated *D. alloeum* gene models DallGr7-9 and four *N. vitripennis* genes NvitGr11-14. Low sequence identity among these genes (15-21%) might suggest an expansion event in a distant wasp ancestor and subsequent loss events in *D. alloeum* and *M. demolitor*. We found 30 GRs in *D. alloeum* grouped with 62 *M. demolitor* genes with indeterminate orthology. Fragmentation of the genome assembly in these regions makes it difficult to use microsynteny analyses to assess orthology and subsequent subfamily expansion in *D. alloeum* and *M. demolitor*. There is strong phylogenetic support for the placement of this group sister to a 32-gene subfamily in *N. vitripennis* (NvitGr15-47), a group containing a single pseudogene in *A. mellifera* (AmelGr11PSE) and is consistent with observations of Robertson *et al*. (2010) suggesting loss of these GR lineages in honeybee. The orthologous group containing *D. alloeum* and *M. demolitor* genes are members of clusters contributing to the enrichment of gustatory-related GO terms in the OrthoVenn analysis. However, as stated above, our annotation of many new GR models in *D. alloeum* permits a more robust view of GR family expansion in braconid wasp ancestors and within *Diachasma* and *Microplitis* lineages. There is also evidence for GR lineage loss in *D. alloeum*. An ortholog group with *N. vitripennis* (NvitGr48-58), *A. mellifera* (AmelGr7, AmelGr12, AmelGrX-ZPSE), and *M. demolitor* (MdemGr51-58, MdemGr72) genes had no apparent ortholog in *D. alloeum*.

Overall, our phylogenetic reconstruction suggests that many GR lineages are the result of extensive gene duplications and differentiation following divergence from an ancestral parasitoid wasp (*Diachasma*-*Microplitis*-*Nasonia* ancestor) and from an ancestral braconid wasp (*Diachasma*-*Microplitis* ancestor). The low sequence similarity among GR sequences within and between species makes it difficult to resolve orthologous relationships among these genes and the timing of duplication/loss events.

We identified a total of 56 ionotropic receptors in *D. alloeum*, representing a considerable expansion relative to *A. mellifera* and *M. mediator* (Table 3). We found 15 “antennal IRs” contained in ortholog groups and 41 “divergent IRs” specific to *D. alloeum*. We confirmed gene predictions for 12 IRs, changed 12 models, and generated 32 new gene annotations. 37 of 56 IR genes (~66%) were arranged in tandem arrays, including repeated antennal IRs (*e.g.* DallIr64a-a.6, Dall75u-u.3) and divergent IRs (*e.g.* DallIr101-108, DallIr109-114). All annotated IRs, with the exception of DallIr124-141, contain at least one of the ligand-gated ion channel domain (PF10613 and/or PF00060). Most IRs, with the exception of members of basal IRs DallIR8a and DallIR25a, lack the characteristic ATP domains (PF01094) of iGLURs (Croset *et al*., 2010).

Phylogenetic analysis of IR sequences shows strong bootstrap support for all antennal IRs within their corresponding ortholog groups (Figure 2). Our phylogeny supports the placement of single-copy DallIr8a, DallIr68a, DallIr76b, and DallIr93a in 1:1 ortholog groups. DallIr25a groups with Ir25a orthologs from other species, which is duplicated in *M. mediator*. DallIr21a groups with orthologs in *N. vitripennis* and *M. mediator*, while this gene has apparently been lost in *A. mellifera*. We recovered six genes in the DallIr64a clade with three orthologs from *N. vitripennis* and two orthologs from *M. mediator*. The three DallIr75u genes group with two orthologs from *N. vitripennis* and one each from *A. mellifera* and *M. mediator.* The divergent IR genes show more complicated patterns of gene duplication and loss, similar to those of the ORs and GRs. Of the 41 divergent IRs, 31 are contained in two large groups; an 18-gene grouping with a single *M. mediator* ortholog (DallIr101-118) and a 13-gene grouping with a large *N. vitripennis* clade (DallIr124-136P+F). These subfamilies may have independently expanded in *Diachasma*/*Nasonia* or alternatively were lost in *Microplitis*. DallIr119, DallIr120, and DallIr121 each form groups with orthologs from other parasitic wasps, and DallIr122/123 represent a recent duplication event specific to the *Diachasma* lineage. DallIr137PSE-140 are nested within a large *N. vitripennis* IR expansion with indeterminate orthology.

We found 15 odorant binding proteins (OBPs) in the *D. alloeum* genome (Table 3). Gene content for this family is relatively consistent among hymenopteran insects studied here, with the exception of the substantial expansion of 98 OBPs in *N. vitripennis* (Vieira *et al*., 2012). OBPs are highly divergent, with pairwise comparisons between *D. alloeum* amino acid sequences having as low as 8% identity. Additionally, there is high interspecies variability across OBPs, with pairwise comparisons in the 3-99% identity range. Despite extensive sequence variability, all *D. alloeum* OBPs contain InterPro hits representative of this family (IPR006170/IPR036728). In addition, OBPs are readily identifiable by the presence of six well-conserved cysteines that form three disulfide bridges as a major structural component of these proteins (Tegoni *et al*., 2004, Pelosi *et al*., 2014). All annotated OBPs in *D. alloeum* contain the canonical six-cysteine structure. Nine DallObps are in tandem repeats, however the notable number of single genes and tandem repeats near edges of scaffolds prevents a comprehensive evaluation of this pattern in the *D. alloeum* genome.

The phylogenetic relationships of OBPs in hymenopterans investigated in this study are shown in Supplementary Figure S8. We recovered DallObp1 and DallObp2 in two separate ortholog groups containing a single gene from each species. Aside from some species-specific OBP expansions, phylogenetic support for ortholog groupings is generally poor. Although these genes are broadly conserved across insect species and have demonstrable function in insect olfaction, the complicated patterns of gene duplication and loss and fragmentation of these genomes makes it difficult for us to infer orthology among surveyed OBPs.

We identified nine chemosensory proteins (CSPs) in *D. alloeum*, similar to the number of CSPs found in other hymenopteran insects (Table 3). We confirmed all CSP gene predictions made by NCBI, and no models were newly generated. Consistent with observations in Forêt *et al*., (2007) regarding tandem arrangement of CSPs in model insect species, five of nine genes were within 50,000 bp of another CSP, with one additional gene near the edge of a small scaffold. All CSP genes contain an InterPro domain specific to this gene family (IPR005055). CSPs are more conserved relative to other chemosensory gene families, with pairwise sequence identity percentages ranging from 15-58% among *D. alloeum* homologs. In addition, all *D. alloeum* CSPs possess four conserved cysteine residues that form two disulfide bridges in these proteins (Tegoni *et al*., 2004).

The phylogeny of CSPs in select hymenopteran insects is shown in Supplementary Figure S9. We recovered no monophyletic groups containing a single gene from each insect, suggesting this family has experienced patterns of lineage-specific gene duplication and/or loss. The interspecific genetic distances are lower for CSPs than other chemosensory families, and pairwise identity values range from 11-68%. Overall, this family has experienced more consistent gene numbers and broader conservation across hymenopteran species.

In summary, this gene set is an important resource for future studies of the evolutionary history of *Diachasma* chemosensory genes. First, it is critical to ascertain the members of the *D. alloeum* chemosensory repertoire that operate specifically in chemosensory behavior. While the families are generally well conserved across insects, the challenge of orthology assessment and the limited functional study of these genes makes it difficult to estimate the precise chemosensory inventory of *D. alloeum*. ORs operate specifically in odorant recognition, and the expansion of OR genes in insects may have been adaptive during the transition to terrestrial life (Robertson *et al*., 2003, but see Missbach *et al*., 2014). Although relatively understudied, the IR family has a likely protostome origin, and conservation of multiple orthologs initially identified in *D. melanogaster* suggest an important function of IR genes in olfaction across insects (Rytz *et al*., 2013). Conversely, the origin of GRs dates back to the Placozoa, and GR-like genes in basal animals function in development, not chemosensation (Robertson, 2015, Saina *et al*., 2015). The OBP and CSP transporter families have roles in chemical ligand delivery to chemosensory receptors but also function in release of pheromones, reproductive processes, and embryonic development (Pelosi *et al*., 2018). Transcriptome datasets used for *D. alloeum* gene predictions were taken from pooled whole male and female wasps, so we cannot exclude the possibility that some genes have non-chemosensory roles. Future studies should incorporate tissue-specific RNA datasets to provide stronger support for genetic components of chemosensation in *D. alloeum*.

Second, chemosensory genes are candidates for differential selective regimes in apple and hawthorn populations of *D. alloeum*. *Rhagoletis pomonella* host flies use olfactory cues from ripening fruit to identify suitable sites for mating and oviposition (Linn *et al*., 2003). Like *R. pomonella*, *D. alloeum* parasitoids have demonstrated odor preferences for their host fruits, representing a potential prezygotic reproductive barrier preventing mating between wasp populations utilizing different hosts (Forbes *et al*., 2009). Evolutionary rate and differential expression analyses of chemosensory genes in *D. alloeum* populations could be potential areas of inquiry.

Third, chemosensory gene evolution could be influenced by transitions in reproductive strategies in *Diachasma*. Wasp courtship is mediated by the male perception of sex pheromones produced by females (Boush & Baerwald, 1967). Across arthropods, chemosensory genes demonstrate differential expression in males and females (*e.g.* Zhou *et al*., 2012, Shiao *et al*., 2013, Eyun *et al*., 2017). Chemosensory genes showing strong sex bias may be candidates for degradation in an asexual genome, such as those involved in female signaling or male recognition of mate signals (Normark *et al*., 2003; Tabata *et al*., 2017). Future studies could assess sex-specific expression of chemosensory genes in *D. alloeum* and corresponding evolutionary patterns in its asexual relative *D. muliebre*.

### References

Bao Z, Eddy SR. 2002. Automated de novo identification of repeat sequence families in	sequenced genomes. Genome Res 12:1269-1276. [doi.org/10.1101%2Fgr.88502](https://dx.doi.org/10.1101%2Fgr.88502)

Bao W, Kojima KK, Kohany O. 2015. Repbase Update, a database of repetitive elements in	eukaryotic genomes. Mobile DNA-UK 6:11. [doi.org/10.1186/s13100-015-0041-9](https://doi.org/10.1186/s13100-015-0041-9)

Benton R, Vannice KS, Gomez-Diaz C, Vosshall LB. 2009. Variant ionotropic glutamate	receptors as chemosensory receptors in *Drosophila*. Cell 136:149-162.	[doi.org/10.1016/j.cell.2008.12.001](https://doi.org/10.1016/j.cell.2008.12.001)

Bernt M, *et al*. 2013. MITOS: improved de novo metazoan mitochondrial genome annotation.	Mol Phylogenet Evol 69:313-319. [doi.org/10.1016/j.ympev.2012.08.023](https://doi.org/10.1016/j.ympev.2012.08.023)

Birney E, Clamp M, Durbin R. 2004. GeneWise and genomewise. Genome Res 14:988-995.	[doi.org/10.1101/gr.1865504](https://genome.cshlp.org/content/14/5/988.short)

Boush GM, Baerwald RJ. 1967. Courtship behavior and evidence for a sex pheromone in the	apple maggot parasite, *Opius alloeus* (Hymenoptera: Braconidae). Ann Entomol Soc Am	60:865-866.

Burke GR, Walden KKO, Whitfield JB, Robertson HM, Strand MR. 2014. Widespread genome	reorganization of an obligate virus mutualist. PLOS Genet 10:e1004660.	[doi.org/10.1371/journal.pgen.1004660](https://doi.org/10.1371/journal.pgen.1004660)

Capella-Gutiérrez S, Silla-Martínez JM, Gabaldón T. 2009. trimAl: a tool for automated	alignment trimming in large-scale phylogenetic analyses. Bioinformatics 25:1972-1973.	<https://doi.org/10.1093/bioinformatics/btp348>

Dierckxsens N, Mardulyn P, Smits G. 2016. NOVOPlasty: de novo assembly of organelle	genomes from whole genome data. Nucleic Acids Res 45:e18-e18.	[doi.org/10.1093/nar/gkw955](https://doi.org/10.1093/nar/gkw955)

Edgar RC. 2004. MUSCLE: multiple sequence alignment with high accuracy and high	throughput. Nucleic Acids Res 32:1792-1797. [doi.org/10.1093/nar/gkh340](https://doi.org/10.1093/nar/gkh340)

English AC, *et al*. 2012. Mind the gap: upgrading genomes with Pacific Biosciences RS long-	read sequencing technology. PLOS One 7:e47768. [doi.org/10.1371/journal.pone.0047768](https://doi.org/10.1371/journal.pone.0047768)

Eyun S, *et al*. 2017. Evolutionary history of chemosensory-related gene families across the	Arthropoda. Mol Biol Evol 34:1838-1862. <https://doi.org/10.1093/molbev/msx147>

Finn RD, *et al*. 2016. The Pfam protein families database: Towards a more sustainable future.	Nucleic Acids Res 44, 279-285. [doi.org/10.1093/nar/gkv1344](https://doi.org/10.1093/nar/gkv1344)

Forbes AA, Powell THQ, Stelinski LL, Smith JJ, Feder JL. 2009. Sequential sympatric	speciation across trophic levels. Science 323:776-779. doi.org/10.1126/science.1166981

Forbes AA, Rice LA, Stewart NB, Yee WL, Neiman M. 2013. Niche differentiation and	colonization of a novel environment by an asexual parasitic wasp. J Evol Biol 26:1330-	1340. [doi.org/10.1111/jeb.12135](https://doi.org/10.1111/jeb.12135)

Forêt S, Maleszka R. 2006. Function and evolution of a gene family encoding odorant binding-	like proteins in a social insect, the honey bee (*Apis mellifera*). Genome Res 16:1404-	1413. [doi.org/10.1101/gr.5075706](https://genome.cshlp.org/content/early/2006/10/25/gr.5075706.short)

Forêt S, Wanner KW, Maleszka R. 2007. Chemosensory proteins in the honey bee: Insights from	the annotated genome, comparative analyses and expressional profiling. Insect Biochem	Molec 37:19-28. [doi.org/10.1016/j.ibmb.2006.09.009](https://doi.org/10.1016/j.ibmb.2006.09.009)

Gibson JD, Niehuis O, Verrelli BC, Gadau J. 2010. Contrasting patterns of selective constraints	in nuclear-encoded genes of the oxidative phosphorylation pathway in holometabolous	insects and their possible role in hybrid breakdown in *Nasonia*. Heredity 104:310-317.	[doi.org/10.1038/hdy.2009.172](https://www.nature.com/articles/hdy2009172)

Grabherr MG, *et al*. 2011. Full-length transcriptome assembly from RNA-Seq data without a	reference genome. Nat Biotechnol 29:644-652. doi.org/10.1038/nbt.1883

Haas BJ, *et al*. 2013. De novo transcript sequence reconstruction from RNA-seq using the	Trinity platform for reference generation and analysis. Nature Protoc 8:1494-1512.	doi.org/10.1038/nprot.2013.084

Hilgenboecker K, Hammerstein P, Schlattmann P, Telschow A, Werren JH. 2008. How many	species are infected with *Wolbachia*? – a statistical analysis of current data. FEMS	Microbiol Lett 281:215-220. [doi.org/10.1111/j.1574-6968.2008.01110.x](https://doi.org/10.1111/j.1574-6968.2008.01110.x)

Hoede C, *et al*. 2014. PASTEC: An automatic transposable element classification tool. PLoS	ONE. 9, e91929. [doi.org/10.1371/journal.pone.0091929](https://doi.org/10.1371/journal.pone.0091929)

Hotopp JCD, *et al*. 2007. Widespread lateral gene transfer from intracellular bacteria to	multicellular eukaryotes. Science 317:1753-1756. [doi.org/10.1126/science.1142490](http://science.sciencemag.org/content/317/5845/1753)

Kearse M, *et al*. 2012. Geneious Basic: an integrated and extendable desktop software platform	for the organization and analysis of sequence data. Bioinformatics 28:1647-1649.	<https://doi.org/10.1093/bioinformatics/bts199>

Kelley DR, Schatz MC, Salzberg SL. 2010. Quake: quality-aware detection and correction of	sequencing errors. Genome Biol 11:R116. <https://doi.org/10.1186/gb-2010-11-11-r116>

Kohany O, Gentles AJ, Hankus L, Jurka J. 2006. Annotation, submission and screening of	repetitive elements in Repbase: RepbaseSubmitter and Censor. *BMC Bioinformatics*	7:474. [doi.org/10.1186/1471-2105-7-474](https://doi.org/10.1186/1471-2105-7-474)

Krieger J, Klink O, Mohl C, Raming K, Breer H. 2003. A candidate olfactory receptor subtype	highly conserved across different insect orders. J Comp Physiol A 189:519-526.	[doi.org/10.1007/s00359-003-0427-x](https://link.springer.com/article/10.1007/s00359-003-0427-x)

Laetsch DR, Blaxter ML. 2017. BlobTools: Interrogation of genome assemblies. F1000Research	6:1287. doi.org/10.12688/f1000research.12232.1

Li H. 2013. Aligning sequence reads, clone sequences and assembly contigs with BWA-MEM.	arXiv:1303.3997.

Li Y, *et al*. 2017. The molecular evolutionary dynamics of oxidative phosphorylation	(OXPHOS) genes in Hymenoptera. BMC Evol Biol 17:269.	<https://doi.org/10.1186/s12862-017-1111-z>

Linn C, *et al*. 2003. Fruit odor discrimination and sympatric host race formation in *Rhagoletis*.	Proc Nat Acad Sci USA 100:11490-11493. <https://doi.org/10.1073/pnas.1635049100>

Luo R, *et al*. 2012. SOAPdenovo2: an empirically improved memory-efficient short-read *de	novo* assembler. Gigascience 1:18. [doi.org/10.1186/2047-217X-1-18](https://doi.org/10.1186/2047-217X-1-18)

Missbach C, *et al*. 2014. Evolution of insect olfactory receptors. Elife 3:e02115.	[doi.org/10.7554/eLife.02115](https://doi.org/10.7554/eLife.02115)

Neiman M, Hehman G, Miller JT, Logsdon Jr JM, Taylor DR. 2009. Accelerated mutation	accumulation in asexual lineages of a freshwater snail. Mol Biol Evol 27:954-963.	<https://doi.org/10.1093/molbev/msp300>

Normark BB, Judson OP, Moran NA. 2003. Genomic signatures of ancient asexual lineages.	Biol J Linn Soc 79:69-84. <https://doi.org/10.1046/j.1095-8312.2003.00182.x>

Normark BB, Moran NA. 2000. Testing for the accumulation of deleterious mutations in asexual	eukaryote genomes using molecular sequences. J Nat Hist 34:1719-1729.	<https://doi.org/10.1080/00222930050122147>

Oliveira DCSG, *et al*. 2009. Identification and characterization of the *doublesex* gene of	*Nasonia*. Insect Mol Biol 18:315-324. [doi.org/10.1111/j.1365-2583.2009.00874.x](https://doi.org/10.1111/j.1365-2583.2009.00874.x)

Park D, *et al*. 2015. Uncovering the novel characteristics of Asian honey bee, *Apis cerana*, by	whole genome sequencing. BMC Genomics 16:1. [doi.org/10.1186/1471-2164-16-1](https://doi.org/10.1186/1471-2164-16-1)

Pascar J, Chandler CH. 2018. A bioinformatics approach to identifying *Wolbachia* infections in	arthropods. PeerJ 6:e5486. [doi.org/10.7717/peerj.5486](https://peerj.com/articles/5486/)

Pelosi P, Mastrogiacomo R, Iovinella I, Tuccori E, Persaud KC. 2014. Structure and	biotechnological applications of odorant-binding proteins. Appl Microbiol Biot 98:61-70.	<https://doi.org/10.1007/s00253-013-5383-y>

Pelosi P, Iovinella I, Zhu J, Wang G, Dani FR. 2018. Beyond chemoreception: diverse tasks of	soluble olfactory proteins in insects. Biol Rev 93:184-200.	<https://doi.org/10.1111/brv.12339>

 Porcelli D, Barsanti P, Pesole G, Caggese C. 2007. The nuclear OXPHOS genes in insecta: a	common evolutionary origin, a common cis-regulatory motif, a common destiny for gene	duplicates. BMC Evol Biol 7:215. <https://doi.org/10.1186/1471-2148-7-215>

Poynton HC, *et al*. 2018. The toxicogenome of *Hyalella azteca*: a model for sediment	ecotoxicology and evolutionary toxicology. Environ Sci Tech 52:6009-6022.	[doi.org/10.1021/acs.est.8b00837](https://pubs.acs.org/doi/abs/10.1021%2Facs.est.8b00837)

Price AL, Jones NC, Pevzner PA. 2005. De novo identification of repeat families in large	genomes. Bioinformatics 21:i351-i358. [doi.org/10.1101/gr.88502](https://doi.org/10.1101/gr.88502)

Rand DM, Haney RA, Fry AJ. 2004. Cytonuclear coevolution: the genomics of cooperation.	Trends Ecol Evol 19:645-653. <https://doi.org/10.1016/j.tree.2004.10.003>

Robertson HM. 2015. The insect chemoreceptor superfamily is ancient in animals. Chem Senses	40:609-614. <https://doi.org/10.1093/chemse/bjv046>

Robertson HM, Gadau J, Wanner KW. 2010. The insect chemoreceptor superfamily of the	parasitoid jewel wasp *Nasonia vitripennis*. Insect Mol Biol 19:121-136.	<https://doi.org/10.1111/j.1365-2583.2009.00979.x>

Robertson HM, Wanner KW. 2006. The chemoreceptor superfamily in the honey bee, *Apis	mellifera*: expansion of the odorant, but not gustatory, receptor family. Genome Res	16:1395-1403. [doi.org/10.1101/gr.5057506](http://www.genome.org/cgi/doi/10.1101/gr.5057506)

Robertson HM, Warr CG, Carlson JR. 2003. Molecular evolution of the insect chemoreceptor	gene superfamily in *Drosophila melanogaster*. Proc Nat Acad Sci USA 100:14537-	14542. <https://doi.org/10.1073/pnas.2335847100>

Rytz R, Croset V, Benton R. 2013. Ionotropic receptors (IRs): chemosensory ionotropic	glutamate receptors in *Drosophila* and beyond. Insect Biochem Molec 43:888-897.	<https://doi.org/10.1016/j.ibmb.2013.02.007>

Saina M, *et al*. 2015. A cnidarian homologue of an insect gustatory receptor functions in	developmental body patterning. Nature Commun 6:6243.	<http://dx.doi.org/10.1038/ncomms7243>

Sharbrough J, Luse M, Boore JL, Logsdon Jr JM, Neiman M. 2018. Radical amino acid	mutations persist longer in the absence of sex. Evolution 72:808-824.	<https://doi.org/10.1111/evo.13465>

Shiao M, *et al*. 2013. Transcriptional profiling of adult *Drosophila* antennae by high-throughput	sequencing. Zool Stud 52:42. <https://doi.org/10.1186/1810-522X-52-42>

Simão FA, Waterhouse RM, Ioannidis P, Kriventseva EV, Zdobnov EM. 2015. BUSCO:	assessing genome assembly and annotation completeness with single-copy orthologs.	Bioinformatics 31:3210-3212. [doi.org/10.1093/bioinformatics/btv351](https://doi.org/10.1093/bioinformatics/btv351)

Smit AFA, Hubley R, Green P. RepeatMasker Open-3.0. 1996-2010.	<http://www.repeatmasker.org>

Smit AFA, Hubley R. RepeatModeler Open-1.0. 2008-2015. <http://www.repeatmasker.org>

Stamatakis A. 2006. RAxML-VI-HPC: maximum likelihood-based phylogenetic analyses with	thousands of taxa and mixed models. Bioinformatics 22:2688-2690.	<https://doi.org/10.1093/bioinformatics/btl446>

Tabata J, Ichiki RT, Moromizato C, Mori K. 2017. Sex pheromone of a coccoid insect with	sexual and asexual lineages: fate of an ancestrally essential sexual signal in	parthenogenetic females. J R Soc Interface 14:20170027.	<https://doi.org/10.1098/rsif.2017.0027>

Tegoni M, Campanacci V, Cambillau C. 2004. Structural aspects of sexual attraction and	chemical communication in insects. Trends Biochem Sci 29:257-264.	<https://doi.org/10.1016/j.tibs.2004.03.003>

Trifinopoulos J, Nguyen LT, von Haeseler A, Minh BQ. 2016. W-IQ-TREE: a fast online	phylogenetic tool for maximum likelihood analysis. Nucleic Acids Res 44:W232-W235.	<https://doi.org/10.1093/nar/gkw256>

Verhulst EC, van de Zande L, Beukeboom LW. 2010. Insect sex determination: it all evolves	around *transformer*. Curr Opin Genet Dev 20:376-383.	[doi.org/10.1016/j.gde.2010.05.001](https://doi.org/10.1016/j.gde.2010.05.001)

Vieira FG, *et al*. 2012. Unique features of odorant-binding proteins of the parasitoid wasp	*Nasonia vitripennis* revealed by genome annotation and comparative analyses. PLoS One	7:e43034. <https://doi.org/10.1371/journal.pone.0043034>

Wang Y, Coleman-Derr D, Chen G, Gu YQ. 2015. OrthoVenn: a web server for genome wide	comparison and annotation of orthologous clusters across multiple species. Nucleic Acids	Res 43:W78-W84. [doi.org/10.1093/nar/gkv487](https://doi.org/10.1093/nar/gkv487)

Wei S, Shi M, Sharkey MJ, van Achterberg C, Chen X. 2010. Comparative mitogenomics of	Braconidae (Insecta: Hymenoptera) and the phylogenetic utility of mitochondrial	genomes with special reference to Holometabolous insects. BMC Genomics 11:371.	<https://doi.org/10.1186/1471-2164-11-371>

Weinstock GM, *et al*. 2006. Insights into social insects from the genome of the honeybee *Apis	mellifera*. Nature 443: 931-949. doi.org/10.1038/nature05260

Werren JH, *et al*. 2010. Functional and evolutionary insights from the genomes of three	parasitoid *Nasonia* species. Science 327:343-348. doi.org/10.1126/science.1178028

Werren JH. 1997. [Biology of Wolbachia](https://doi.org/10.1146%2Fannurev.ento.42.1.587). Ann Rev Entomol 42:587-609.	[doi.org/10.1146/annurev.ento.42.1.587](https://doi.org/10.1146/annurev.ento.42.1.587)

Wharton RA, Yoder MJ. 2015. Parasitoids of fruit-infesting tephritidae. Available at	[paroffit.org/public/site/paroffit/home](http://paroffit.org/public/site/paroffit/home). Accessed July 2, 2015.

Wheeler D, Redding AJ, Werren JH. 2013. Characterization of an ancient lepidopteran lateral	gene transfer. PLOS One 8:e59262. <https://doi.org/10.1371/journal.pone.0059262>

Wicker T, *et al*. 2007. A unified classification system for eukaryotic transposable elements.	Nat Rev Genet 8:973. [doi.org/10.1038/nrg2165](https://doi.org/10.1038/nrg2165)

Zdobnov EM, Apweiler R. 2001. InterProScan–an integration platform for the signature-	recognition methods in InterPro. *Bioinformatics* 9:847-848.	<https://doi.org/10.1093/bioinformatics/17.9.847>

Zdobnov EM, *et al*. 2016. OrthoDB v9.1: cataloging evolutionary and functional annotations for	animal, fungal, plant, archaeal, bacterial and viral orthologs. Nucleic Acids Res 45:D744-	D749. <https://doi.org/10.1093/nar/gkw1119>

Zhang F, Broughton RE. 2013. Mitochondrial–nuclear interactions: compensatory evolution or	variable functional constraint among vertebrate oxidative phosphorylation genes?	Genome Biol Evol 5:1781-1791. <https://doi.org/10.1093/gbe/evt129>

Zhou X, *et al*. 2015. Chemoreceptor evolution in hymenoptera and its implications for the	evolution of eusociality. Genome Biol Evol 7:2407-2416. [doi.org/10.1093/gbe/evv149](https://doi.org/10.1093/gbe/evv149)

Zhou X *et al*. 2012. Phylogenetic and transcriptomic analysis of chemosensory receptors in a pair	of divergent ant species reveals sex-specific signatures of odor coding. PLOS Genet	8:e1002930. <https://doi.org/10.1371/journal.pgen.1002930>

### Supplementary Tables

**Supplementary Table S1.** Sex determination genes used as queries for BLAST searches.

| **Gene Name** | **Organism** | **Accession** |
| --- | --- | --- |
| *doublesex* (female) | *N. vitripennis* | NP_001155990.1 |
| *doublesex* (male) | *N. vitripennis* | NP_001155989.1 |
| *doublesex* (female) | *A. mellifera* | ABW99105.1 |
| *doublesex* (male) | *A. mellifera* | ABW99102.1 |
| *transformer* | *N. vitripennis* | NP_001128299.1 |
| *feminizer* | *A. mellifera* | NP_001128300.1 |
| *csd* | *A. mellifera* | ABU68670.1 |

**Supplementary Table S2.** Arthropoda BUSCO analysis for four hymenopteran genomes.

|  | | **Arthropoda BUSCOs (1066 total)** | | | |
| --- | --- | --- | --- | --- | --- |
| **Organism** | **Assembly Name** | **Complete**  **single-copy** | **Complete**  **duplicated** | **Fragmented** | **Missing** |
| *Diachasma alloeum* | GCA_001412515.1  (this study) | 1043 | 10 | 7 | 6 |
| *Apis mellifera* | GCA_000002195.1 (Weinstock *et al*., 2006) | 1038 | 6 | 15 | 7 |
| *Nasonia vitripennis* | GCA_000002325.2  (Werren *et al*., 2010) | 1030 | 8 | 15 | 13 |
| *Microplitis demolitor* | GCA_000572035.2  (Burke *et al*., 2014) | 1038 | 19 | 4 | 5 |

**Supplementary Table S3.** Hymenoptera BUSCO analysis for four hymenopteran genomes.

|  | | **Hymenoptera BUSCOs (4415 total)** | | | |
| --- | --- | --- | --- | --- | --- |
| **Organism** | **Assembly Name** | **Complete**  **single-copy** | **Complete**  **duplicated** | **Fragmented** | **Missing** |
| *Diachasma alloeum* | GCA_001412515.1  (this study) | 4132 | 30 | 138 | 115 |
| *Apis mellifera* | GCA_000002195.1 (Weinstock *et al*., 2006) | 4282 | 13 | 58 | 62 |
| *Nasonia vitripennis* | GCA_000002325.2  (Werren *et al*., 2010) | 4094 | 24 | 192 | 105 |
| *Microplitis demolitor* | GCA_000572035.2  (Burke *et al*., 2014) | 4023 | 48 | 172 | 172 |

**Supplementary Table S4.** Summary of transposable elements (TEs) and other repetitive content in *Diachasma alloeum*. Details and abbreviations provided in Supplementary Table S6.

| Class | Order | Superfamily | Families | Copies | bp Masked | % masked |
| --- | --- | --- | --- | --- | --- | --- |
| DNA transposons | TIR | Academ | 1 | 136 | 173573 | 0.05 |
|  |  | CMC | 3 | 799 | 435933 | 0.11 |
|  |  | Kolobok-Hydra | 1 | 141 | 74544 | 0.02 |
|  |  | Merlin | 4 | 319 | 174926 | 0.05 |
|  |  | MuLE | 2 | 200 | 55433 | 0.01 |
|  |  | PIF | 1 | 136 | 70721 | 0.02 |
|  |  | Sola | 5 | 2604 | 1811472 | 0.48 |
|  |  | TcMar | 22 | 8150 | 4679588 | 1.22 |
|  |  | Zator | 1 | 74 | 8362 | 0.00 |
|  |  | hAT | 9 | 2126 | 1189937 | 0.31 |
|  |  | other | 154 (128) | 62853 | 15762986 | 4.10 |
|  |  | **Total TIR** | **205** | **77538** | **24437475** | **6.37** |
|  | Crypton | Crypton | 1 | 147 | 53722 | 0.01 |
|  | Polinton | Maverick | 28 | 12303 | 10729882 | 2.79 |
|  | RC | Helitron | 7 | 2977 | 2139824 | 0.56 |
|  |  | **Total DNA** | **241** | **92965** | **37360903** | **9.73** |
| Retrotransposons | LTR | Copia | 9 | 2822 | 3140105 | 0.82 |
|  |  | Gypsy | 46 | 10173 | 10879077 | 2.83 |
|  |  | Poa | 19 | 5524 | 4206592 | 1.10 |
|  |  | other | 4 | 624 | 265208 | 0.07 |
|  |  | **Total LTR** | **78** | 19143 | 18490982 | 4.82 |
|  | DIRS | DIRS | 3 | 593 | 438874 | 0.11 |
|  | PLE | Penelope | 6 | 2805 | 1268724 | 0.33 |
|  | LINE | CR1 | 2 | 226 | 72862 | 0.02 |
|  |  | Dong-R4 | 2 | 942 | 1181547 | 0.31 |
|  |  | I | 8 | 1222 | 722177 | 0.19 |
|  |  | Jockey | 3 | 626 | 477712 | 0.12 |
|  |  | L1 | 1 | 188 | 194973 | 0.05 |
|  |  | L2 | 3 | 544 | 840422 | 0.22 |
|  |  | LOA | 3 | 1825 | 2190878 | 0.57 |
|  |  | R1 | 4 | 2158 | 3060582 | 0.80 |
|  |  | RTE | 10 | 2094 | 1859848 | 0.48 |
|  |  | **Total LINE** | **36** | 8657 | 9346592 | 2.43 |
|  | SINE |  | **3** | 525 | 124444 | 0.03 |
|  |  | **Total Retro** | **126** | **31723** | **29669616** | **7.72** |
| **Total classified TEs** | |  | **367** | 450855 | 114417735 | **17.87** |
| Unknown | |  | 1048 | 8732 | 439580 | 29.79 |
| Low complexity sequence | |  |  | 8902 | 2146309 | 0.11 |
| Satellite | |  |  | 55478 | 4050536 | 0.56 |
| Simple repeats | |  |  | 450855 | 114417735 | 1.05 |
| **Total repetitive sequence** | |  |  |  | **188084679** | **49.37** |

**Supplementary Table S5.** Summary of transposable elements (TEs) and other repetitive content in Nasonia vitripennis. Details and abbreviations provided in Supplementary Table S6.

| Class | Order | Superfamily | Families | Copies | bp Masked | % masked |
| --- | --- | --- | --- | --- | --- | --- |
| DNA transposons | TIR | Academ | 2 | 109 | 91283 | 0.03 |
|  |  | CMC | 13 | 2228 | 789484 | 0.27 |
|  |  | Kolobok-Hydra | 7 | 1285 | 534771 | 0.18 |
|  |  | MuLE | 4 | 334 | 104974 | 0.04 |
|  |  | PIF | 2 | 431 | 355161 | 0.12 |
|  |  | PiggyBac | 1 | 75 | 24507 | 0.01 |
|  |  | Sola | 1 | 607 | 273914 | 0.09 |
|  |  | TcMar | 7 | 1189 | 419829 | 0.14 |
|  |  | hAT | 3 | 798 | 210410 | 0.07 |
|  |  | other | 62 | 8799 | 2160063 | 0.73 |
|  |  | **Total TIR** | **102** | **15855** | **4964396** | **1.68** |
|  | Crypton | Crypton | 2 | 154 | 50707 | 0.02 |
|  | Polinton | Maverick | 41 | 6386 | 3671249 | 1.25 |
|  | RC | Helitron | 21 | 6592 | 2980101 | 1.01 |
|  |  | **Total DNA** | **166** | **28987** | **11666453** | **3.96** |
| Retrotransposons | LTR | Copia | 15 | 1664 | 1270913 | 0.43 |
|  |  | Gypsy | 75 | 7627 | 6050339 | 2.05 |
|  |  | Poa | 9 | 598 | 596847 | 0.20 |
|  |  | other | 4 | 282 | 488918 | 0.17 |
|  |  | **Total LTR** | **103** | **10171** | **8407017** | **2.85** |
|  | DIRS | DIRS | 1 | 113 | 125846 | 0.04 |
|  | PLE | Penelope | 4 | 650 | 243196 | 0.08 |
|  | LINE | CR1 | 5 | 329 | 172810 | 0.06 |
|  |  | I | 3 | 541 | 364274 | 0.12 |
|  |  | L2 | 23 | 4936 | 3305819 | 1.12 |
|  |  | LOA | 6 | 687 | 526420 | 0.18 |
|  |  | R1 | 18 | 4315 | 3149203 | 1.07 |
|  |  | RTE | 6 | 782 | 875722 | 0.30 |
|  |  | **Total LINE** | **61** | **11590** | **8394248** | **2.85** |
|  | SINE |  | 3 | 274 | 80563 | 0.03 |
|  |  | **Total Retro** | **172** | **22798** | **17250870** | **5.85** |
| **Total classified TEs** | |  | **338** | **51785** | **28917323** | **9.81** |
| Unknown | |  | **582** | 84269 | 26593342 | 9.03 |
| Low complexity sequence | |  |  |  | 408177 | 0.14 |
| Satellite | |  |  |  | 6497083 | 2.21 |
| Simple repeats | |  |  |  | 5359921 | 1.82 |
| **Total repetitive sequence** | |  |  |  | **67775846** | **23.01** |

**Supplementary Table S6.** Summary of transposable elements (TEs) and other repetitive content in Apis mellifera.

| Class | Order | Superfamily | Families | Copies | bp Masked | % masked |
| --- | --- | --- | --- | --- | --- | --- |
| DNA transposons | TIR | TcMar | 13 | 1524 | 588164 | 0.23 |
|  |  | other | 4 (3) | 1506 | 361131 | 0.14 |
|  |  | **Total TIR** | 1**7** | **3030** | **949295** | **0.37** |
|  | Crypton |  | 0 | 0 | 0 | 0.00 |
|  | Polinton |  | 0 | 0 | 0 | 0.00 |
|  | RC |  | 0 | 0 | 0 | 0.00 |
|  |  | **Total DNA** | **17** | **3030** | **949295** | **0.37** |
| Retrotransposons | LTR | Copia | 1 | 84 | 62234 | 0.02 |
|  | DIRS |  | 0 | 0 | 0 | 0.00 |
|  | PLE |  | 0 | 0 | 0 | 0.00 |
|  | LINE |  | 0 | 0 | 0 | 0.00 |
|  | SINE |  | 0 | 0 | 0 | 0.00 |
|  |  | **Total retro** | **1** | **84** | **62234** | **0.02** |
| **Total classified TEs** | |  | **18** | **3114** | **1011529** | **0.39** |
| Unknown | |  | **147** | 28209 | 8536684 | 3.43 |
| Low complexity sequence | |  |  |  | 2609155 | 1.05 |
| Satellite | |  |  |  | 0 | 0.00 |
| Simple repeats | |  |  |  | 10807174 | 4.34 |
| **Total repetitive sequence** | |  |  |  | **22964542** | **9.22** |

Total family numbers, element counts, total bp masked, and percent of the genome masked for each TE superfamily, organized by hierarchy described by Wicker *et al*. (2007). TIR = terminal inverted repeat transposon, LINE and SINEs = long and interspersed nuclear elements, respectively, LTR = long terminal repeat. The “other” TIR and LTR groups are sequence families with structural characteristics of these TE orders lacking sequence similarity to known TE families; in parentheses is the number of “other” TIRs classified as miniature inverted transposable elements (MITEs) by PASTEClassifier.

### Supplementary Figures

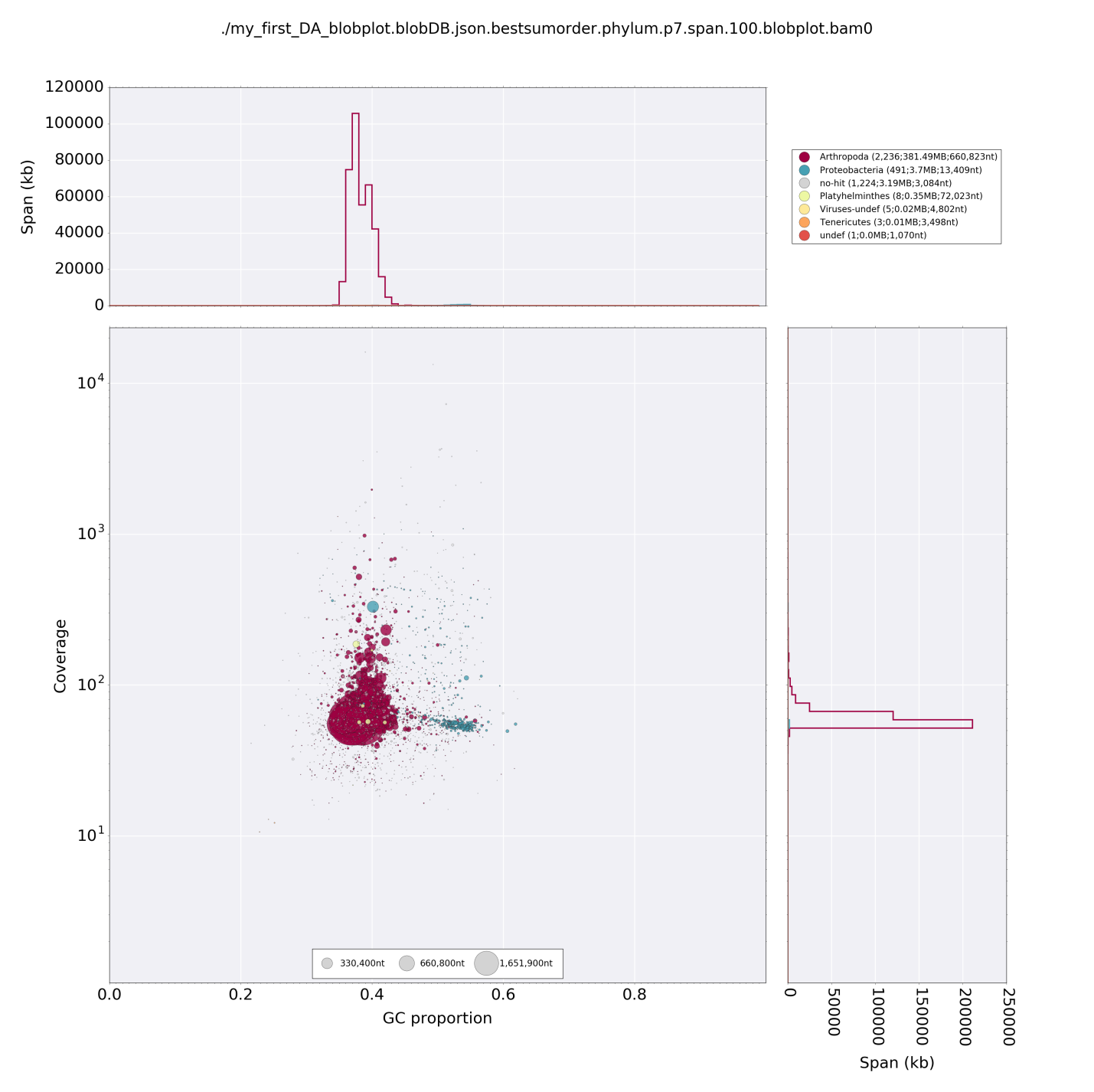

**Supplementary Figure S1.** Blobplot of *Diachasma alloeum* genome assembly. Length, depth of coverage, and GC content are plotted for each scaffold. Dots are color coded according to primary taxonomic identifier using BlobTools ‘taxify.’ Size of dot corresponds to scaffold size. Image generated using BlobTools v1.0 (DOI 10.5281/zenodo.845347, Laetsch & Blaxter, 2017).

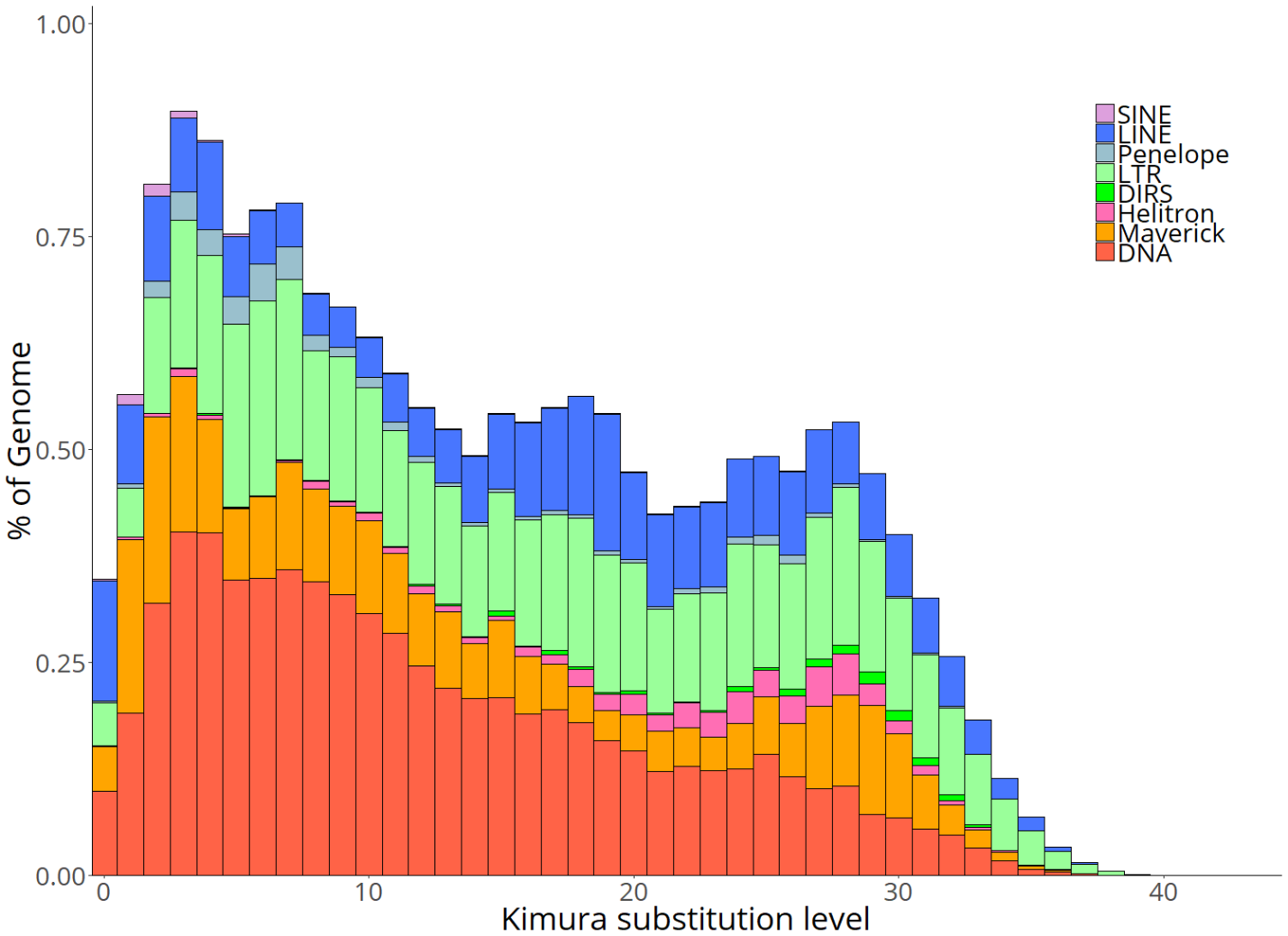

**Supplementary Figure S2.** Repeat landscape of annotated TEs in *Diachasma alloeum*. X axis: Kimura substitution level (CpG adjusted) relative to the family consensus sequence, reflecting the age of the TEs. Younger TEs (more recently inserted into the genome) have lower levels of divergence. Y axis: percent of the genome made up by each TE class for bins of 1 % substitution level.

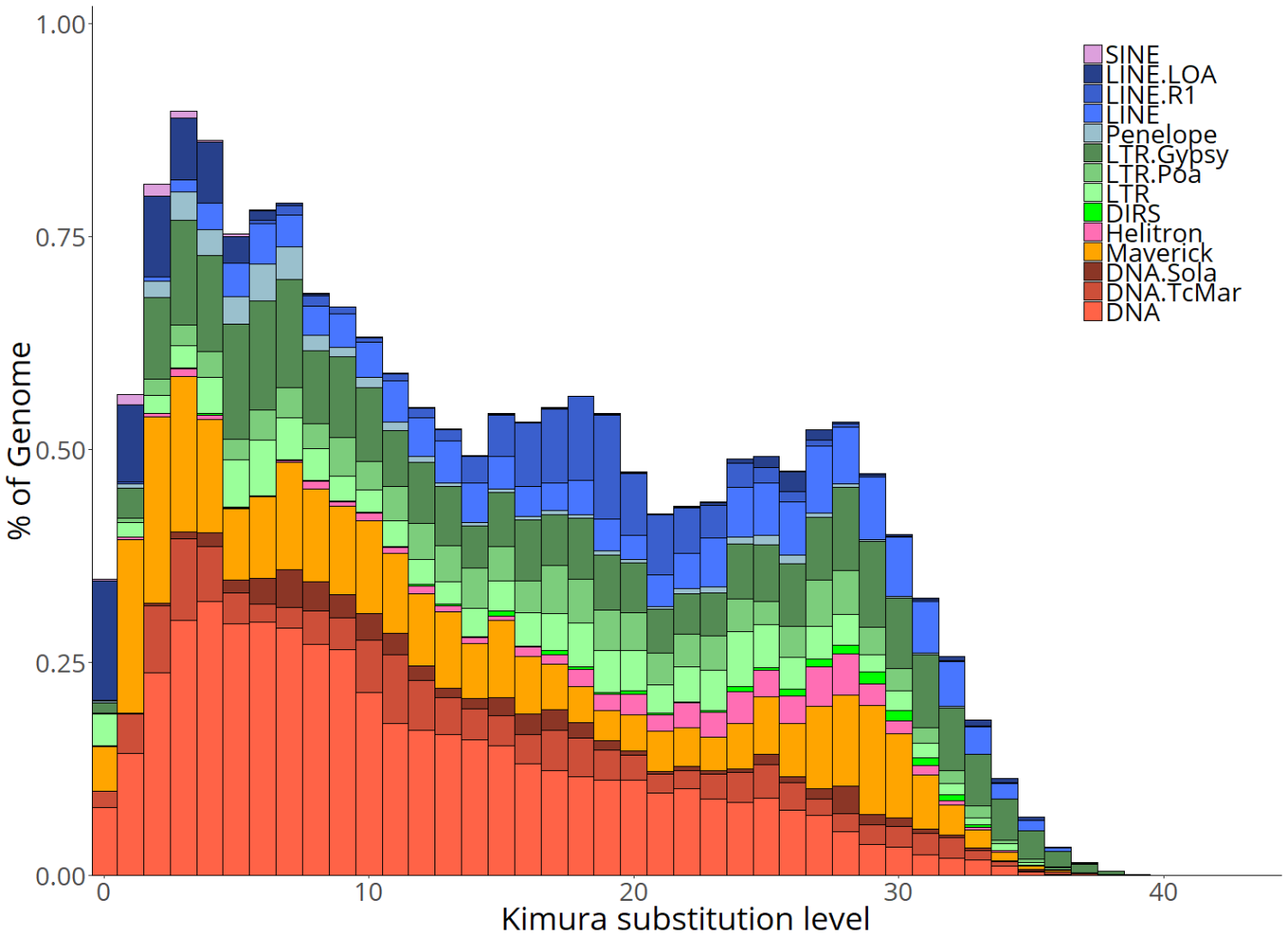

**Supplementary Figure S3.** Repeat landscape of annotated TEs in *Diachasma alloeum*. X axis: Kimura substitution level (CpG adjusted) relative to the family consensus sequence, reflecting the age of the TEs. Younger TEs (more recently inserted into the genome) have lower levels of divergence. Y axis: percent of the genome made up by each TE class for bins of 1 % substitution level. The LTR, LINE, and DNA (referring to TIR) orders contain numerous superfamilies, here the 2 most abundant superfamilies for these groups are included separately from the rest of the masking results.

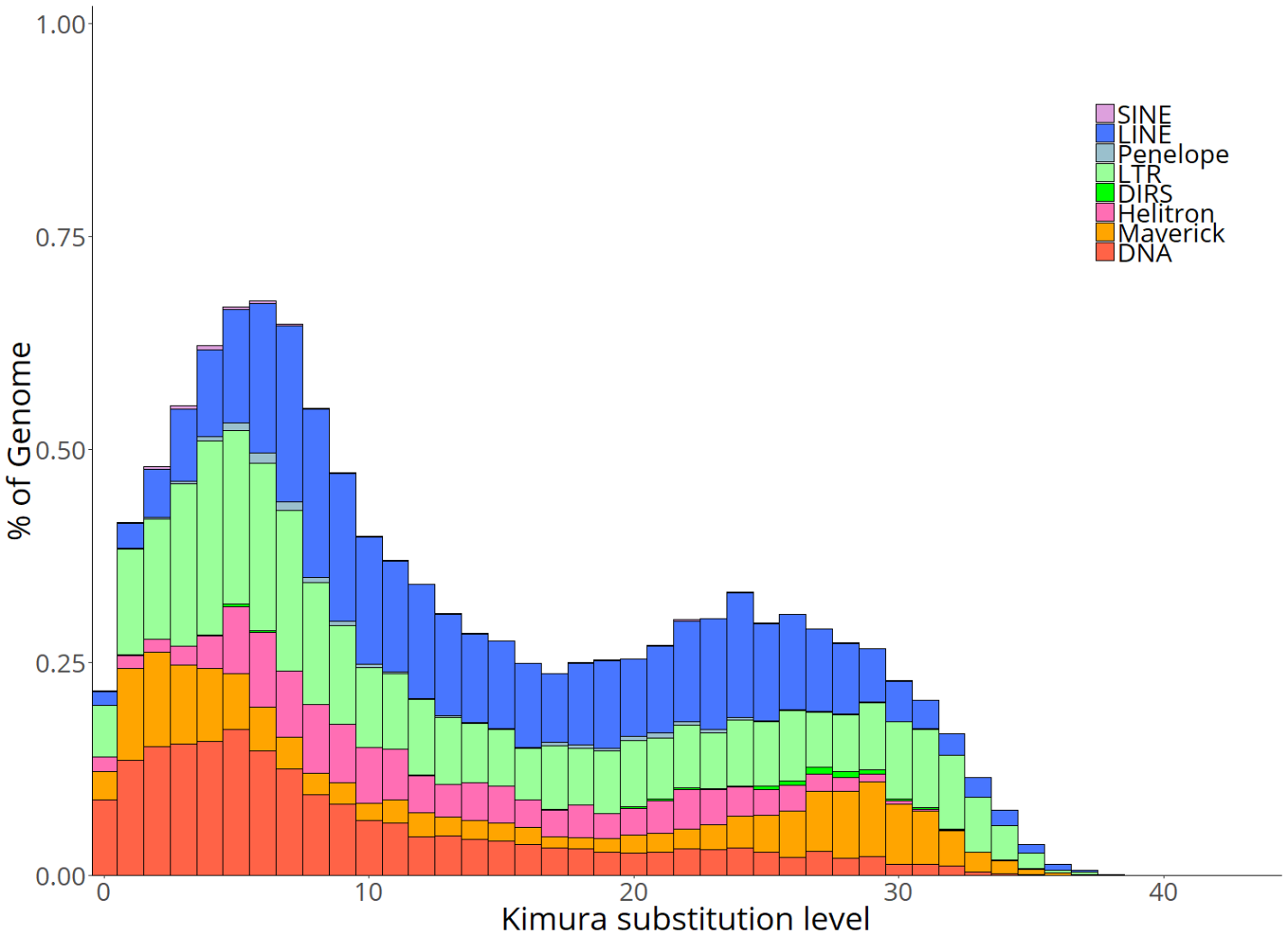

**Supplementary Figure S4.** Repeat landscape of annotated TEs in *Nasoina vitripennis*. X axis: Kimura substitution level (CpG adjusted) relative to the family consensus sequence, reflecting the age of the TEs. Younger TEs (more recently inserted into the genome) have lower levels of divergence. Y axis: percent of the genome made up by each TE class for bins of 1 % substitution level. Y-axis scale is based on the scale fitting for *D. alloeum*.

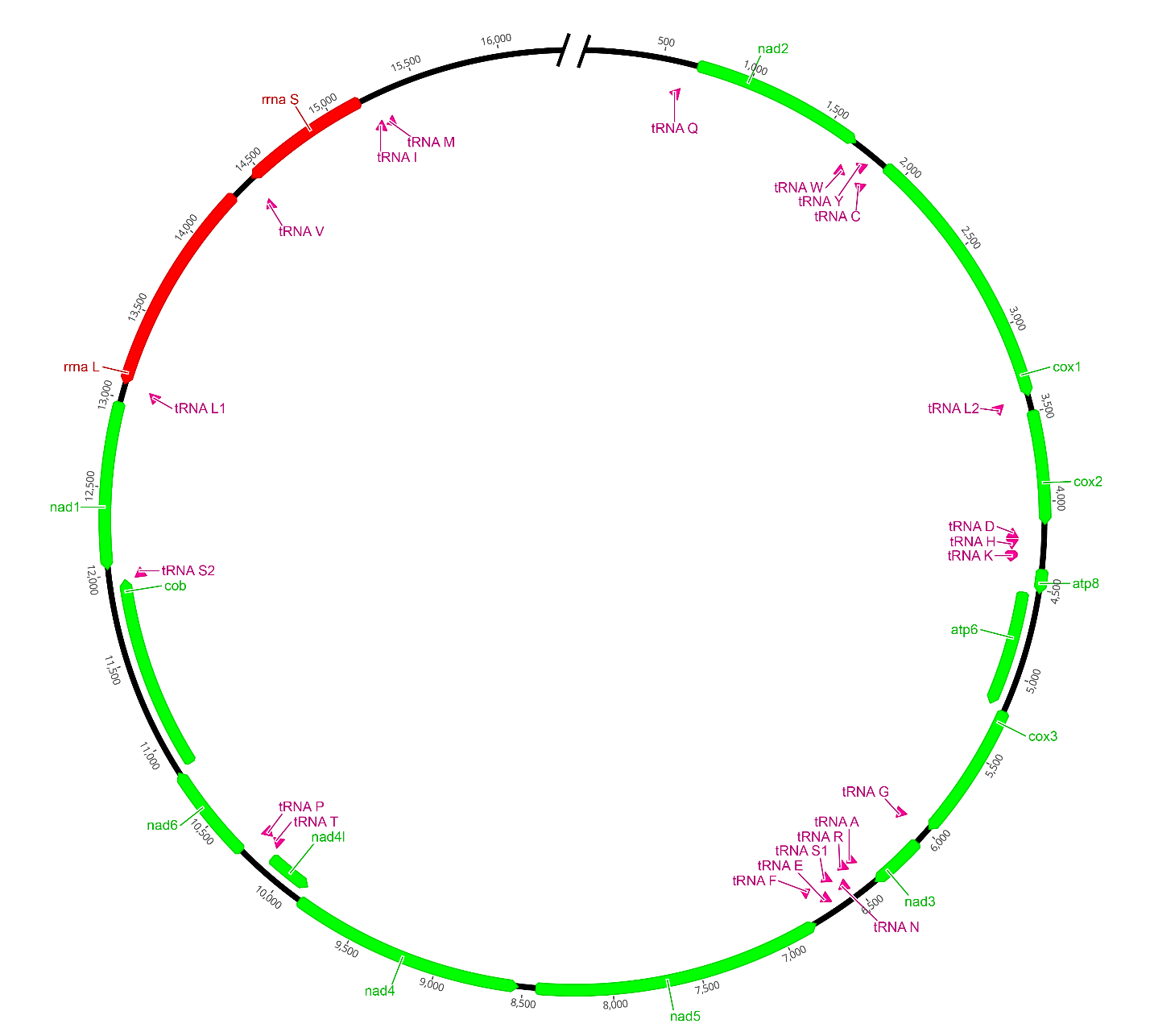

**Supplementary Figure S5.** Diagram of mitochondrial genome of *Diachasma alloeum*. Protein-coding genes are shown in green, rRNAs are shown in red, tRNAs are shown in pink. Leftward and rightward facing arrows indicate direction of transcription. AT-rich control region (top) was not fully assembled by NOVOPlasty. Image generated using Geneious v9.1 (Biomatters, Ltd.).

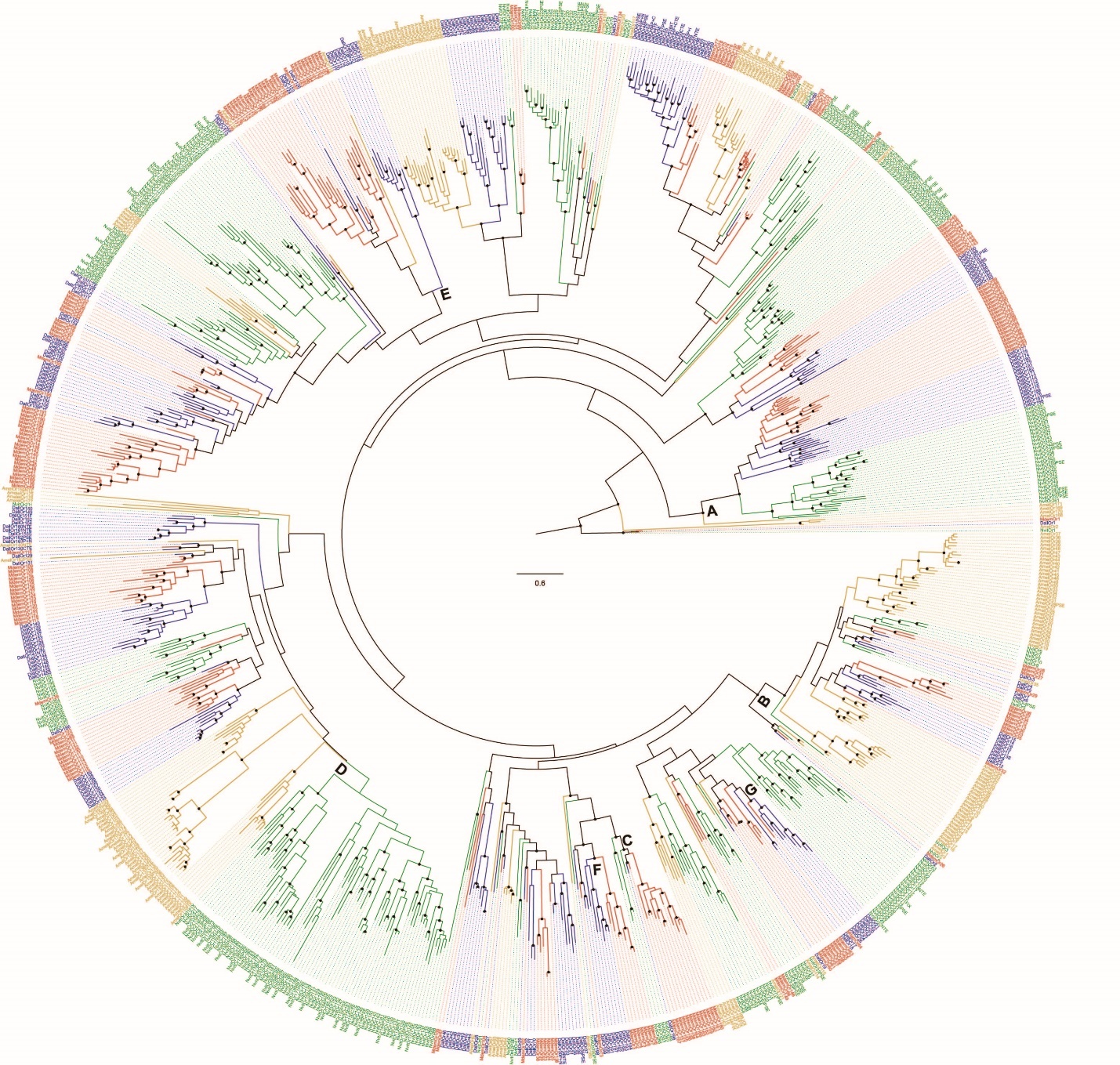

**Supplementary Figure S6.** Phylogenetic tree of ORs from sampled hymenopteran insects.

Taxa surveyed include *Diachasma alloeum* (blue), *Microplitis demolitor* (red), *Nasonia vitripennis* (green), and *Apis mellifera* (orange). Maximum likelihood tree generated using 305 alignment columns. Dots on nodes indicate > 90% bootstrap support. The scale bar indicates the number of amino acid substitutions per site. Letters A-G represent points of interest in the phylogenetic tree.

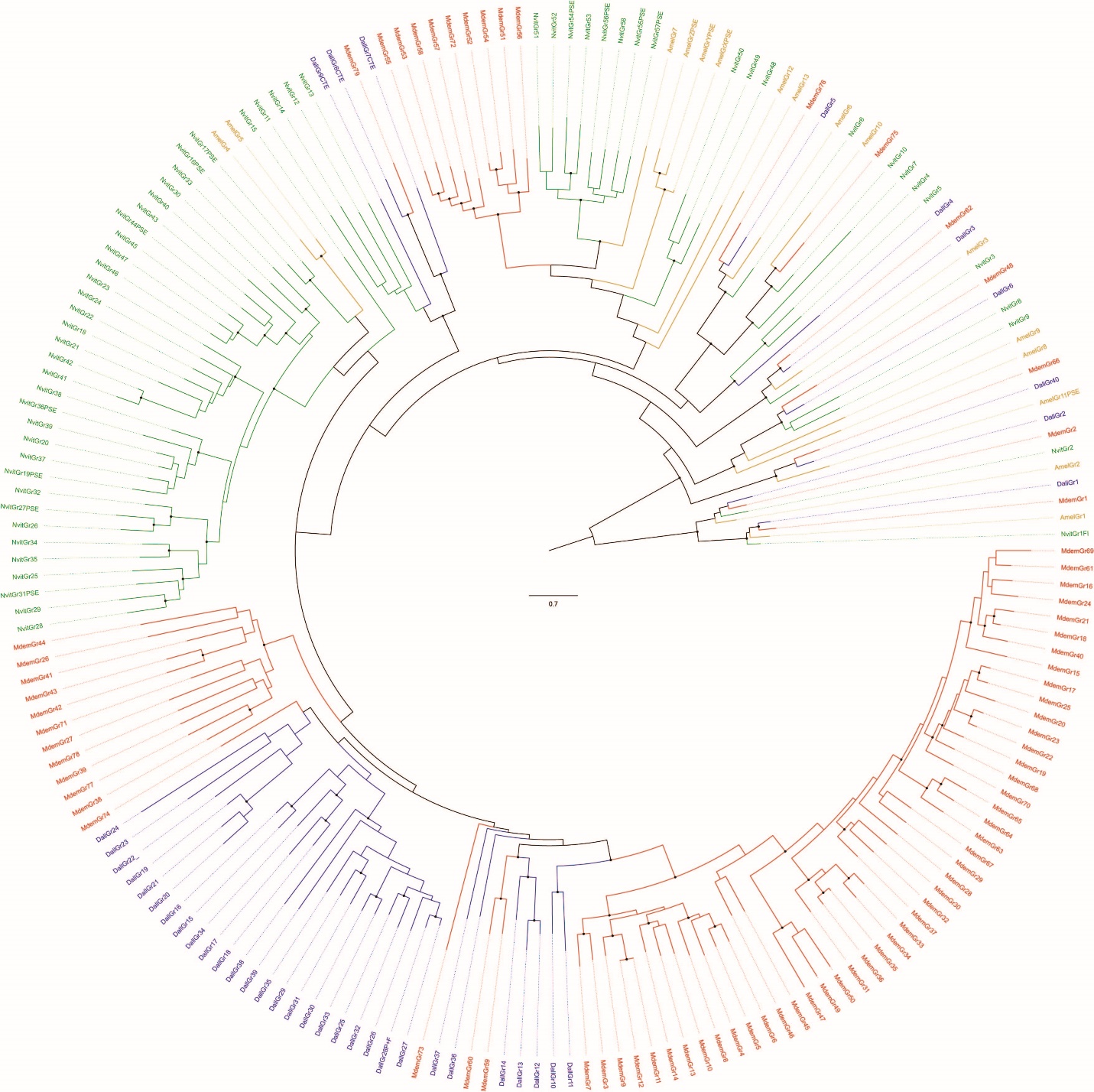

**Supplementary Figure S7.** Phylogenetic subtrees of GRs from sampled hymenopteran insects.

Dall = *Diachasma alloeum*, Mmed = *Microplitis mediator*, Nvit = *Nasonia vitripennis*, Amel = *Apis mellifera*. Maximum likelihood tree generated using 431 alignment columns. Dots on nodes indicate > 90% bootstrap support. The scale bar indicates the number of amino acid substitutions per site.

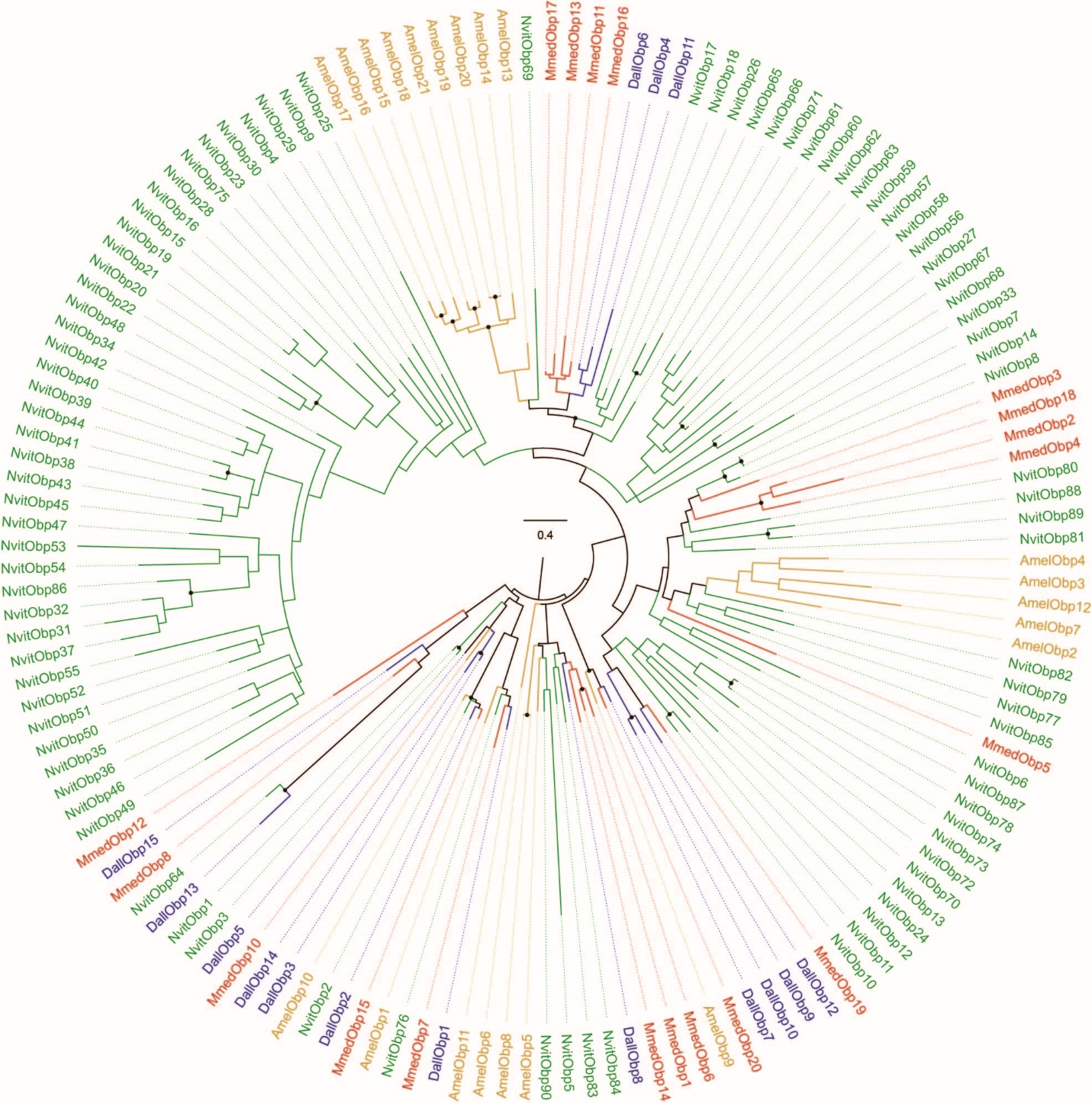

**Supplementary Figure S8.** Phylogenetic subtrees of OBPs from sampled hymenopteran insects.

Dall = *Diachasma alloeum*, Mmed = *Microplitis mediator*, Nvit = *Nasonia vitripennis*, Amel = *Apis mellifera*. Maximum likelihood tree generated using 169 alignment columns. Dots on nodes indicate > 90% bootstrap support. The scale bar indicates the number of amino acid substitutions per site.

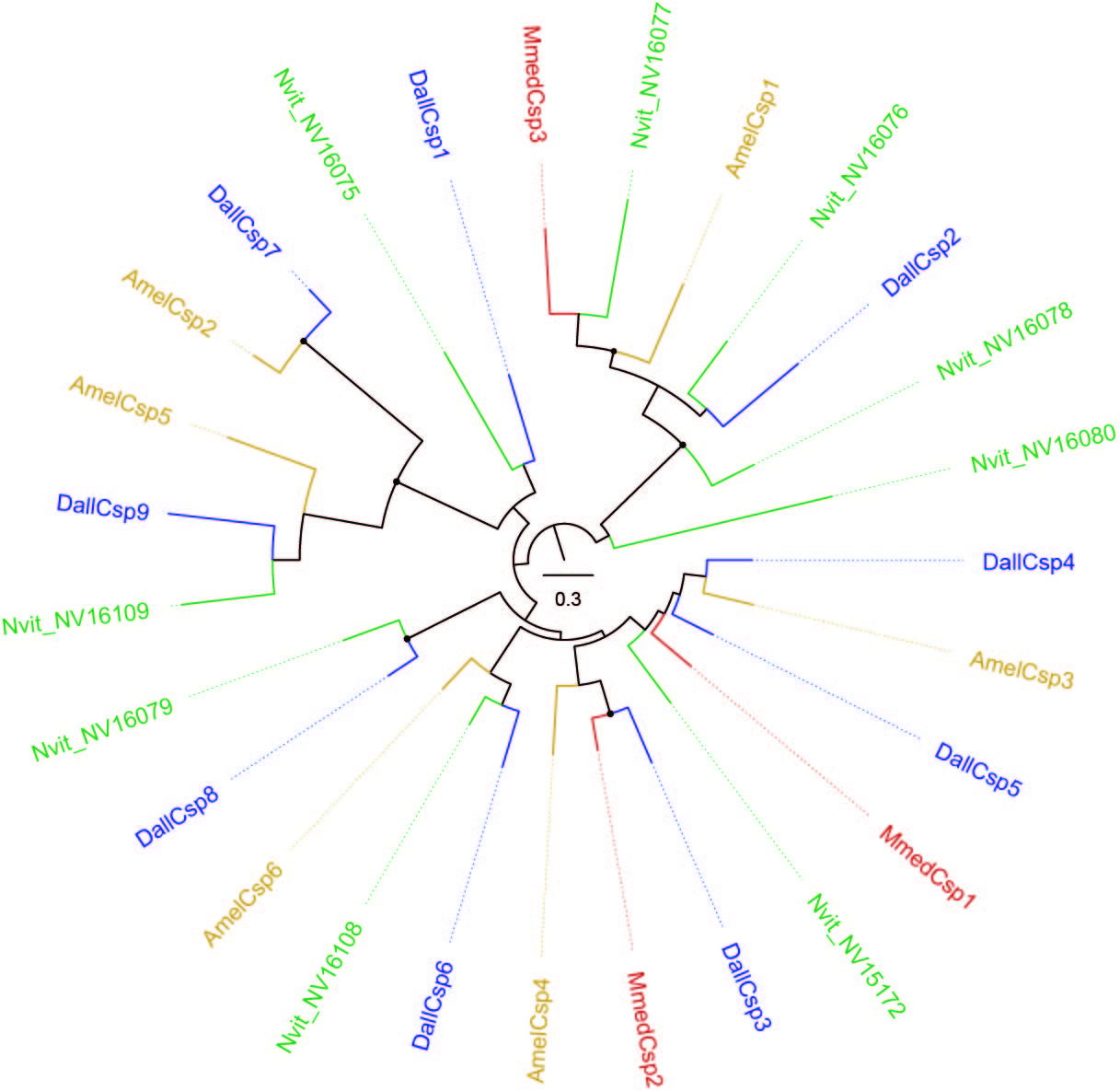

**Supplementary Figure S9.** Phylogenetic subtrees of CSPs from sampled hymenopteran insects. Dall = *Diachasma alloeum*, Mmed = *Microplitis mediator*, Nvit = *Nasonia vitripennis*, Amel = *Apis mellifera*. Maximum likelihood tree generated using 130 alignment columns. Dots on nodes indicate > 90% bootstrap support. The scale bar indicates the number of amino acid substitutions per site.

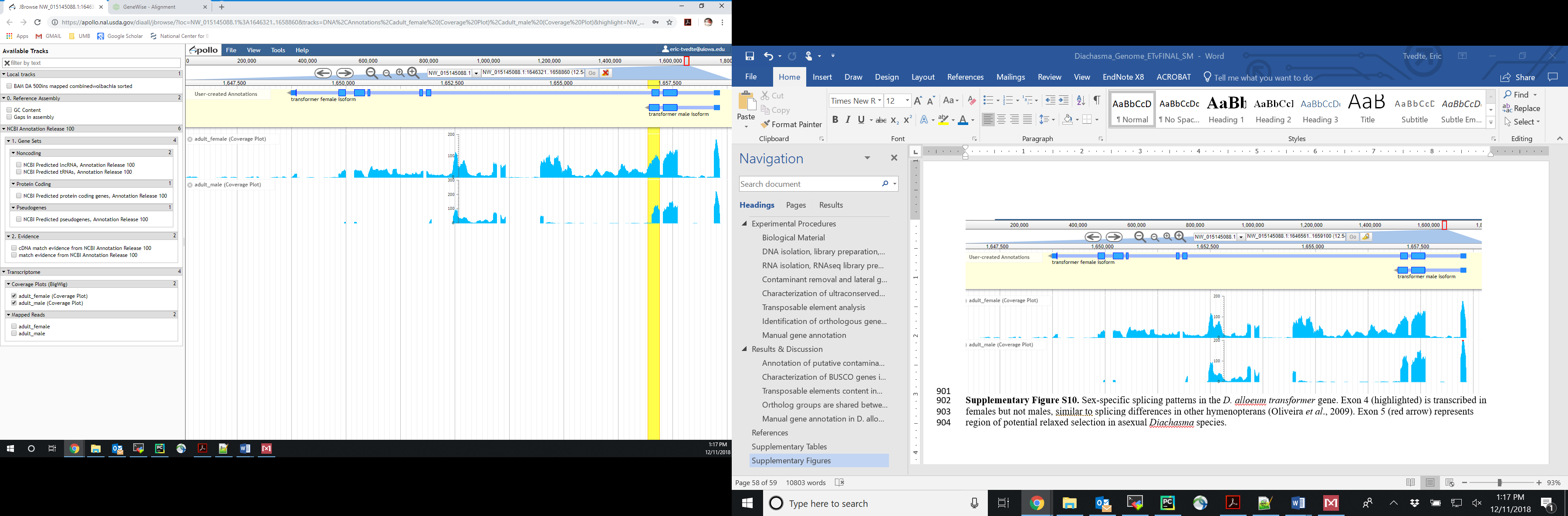

**Supplementary Figure S10.** Sex-specific splicing patterns in the *D. alloeum* *transformer* gene. Predicted sex-specific isoforms are displayed on the upper track, coverage plot of transcriptome datasets are displayed on lower track. Lower transcriptional activity in D. alloeum males could be due to premature stop codon in Exon 2 (highlighted), which is also observed in other insects (Verhulst *et al*., 2010).

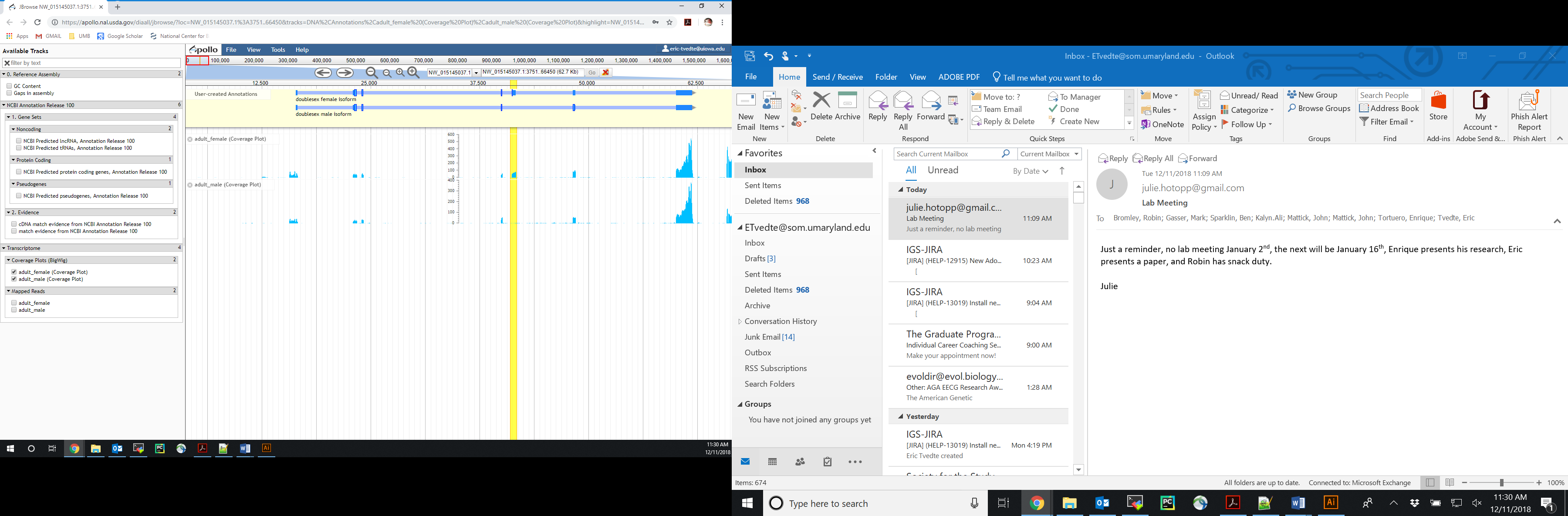

**Supplementary Figure S11.** Sex-specific splicing patterns in the *D. alloeum* *doublesex* gene. Predicted sex-specific isoforms are displayed on the upper track, coverage plot of transcriptome datasets are displayed on lower track. Exon 4 (highlighted) is transcribed in females but not males, similar to splicing differences in other hymenopterans (Oliveira *et al*., 2009). Exon 5 (red arrow) represents region of potential relaxed selection in asexual *Diachasma* species.
